# The chemokine receptor CXCR3 promotes CD8^+^ T cell-dependent lung pathology during influenza pathogenesis

**DOI:** 10.1101/2022.02.14.480379

**Authors:** Kai Guo, Dan J.K Yombo, Jintao Xu, Zhihan Wang, Taylor Schmit, Jitendra Tripathi, Junguk Hur, Jie Sun, Michal A. Olszewski, Nadeem Khan

**Affiliations:** Department of Neurology, University of Michigan, Ann Arbor, MI 48109, USA; Department of Biomedical Sciences, School of Medicine and Health Sciences, University of North Dakota, Grand Forks, ND 58202, USA; Research Service, Ann Arbor VA Health System, Department of Veterans Affairs Health System, Ann Arbor, Michigan, USA; Division of Pulmonary and Critical Care Medicine, Department of Internal Medicine, University of Michigan Health System, Ann Arbor, Michigan, USA; West China School of Basic Medical Sciences & Forensic Medicine, Sichuan University, Chengdu, Sichuan, China; Carter Immunology Center, University of Virginia, Charlottesville, VA, USA; Division of Infectious Diseases and International Health, University of Virginia. Charlottesville, VA, USA; Dept of Oral Biology, College of Dentistry, University of Florida, Gainesville, FL, USA

## Abstract

While the role of CD8^+^ T cells in influenza clearance is established, their contribution to pathological lung injury is increasingly appreciated. To explore if protective versus pathological functions can be linked to CD8^+^ T cell subpopulations, we dissected their responses in influenza-infected murine lungs. Our single-cell RNASeq (scRNAseq) analysis revealed significant diversity in CD8^+^ T cell subpopulations during peak viral load vs. infection-resolved state. While enrichment of Cxcr3^hi^ CD8^+^ T effector (T_eff_) subset was associated with a more robust cytotoxic response, both CD8^+^ T_eff_ and CD8^+^ T central memory (T_CM_) exhibited equally potent effector potential. The scRNAseq analysis identified unique regulons regulating the cytotoxic response in CD8^+^ T cells. The neutralization of CXCR3 mitigated lung injury without affecting viral clearance. IFN-γ was dispensable to regulate the cytotoxic response of Cxcr3^hi^ CD8^+^ T cells. Collectively, our data imply that CXCR3 interception could have a therapeutic effect in preventing influenza-linked lung injury.

**TEASER:** The CXCR3 expressing CD8+ T cell subset causes severe lung pathology and exacerbates disease severity without affecting viral clearance during influenza infection

## INTRODUCTION

Influenza exhibits a complex disease phenotype, ranging from a self-limiting mild infection to severe life-threatening pneumonia. Seasonal influenza vaccines offer limited efficacy (*1*), and influenza remains a significant public health problem with > 30,000 annual deaths and over $10.4 billion healthcare costs (*2*). Severe influenza pneumonia manifestations are profound airway lung and vascular injuries, impacting gas exchange and requiring hospitalization (*3–5*). The recovery is often slow and may leave patients with permanent lung damage (*6–8*). Most of these complications are attributed to host’s own defense mechanisms because host responses while executing control of the viral load drive collateral lung damage (*9, 10*). Viral load in the lung is one of the factors that affect the balance between infection resolution and immune-mediated lung pathology, but paradoxically the acute injury continues and persists in the lungs after the viral clearance (*11, 12*). Despite numerous studies exploring the pathogenesis of influenza, there is a significant knowledge gap in how the host response to influenza turns pathologic and which subsets of the immune cells are the major drivers of pathology during and post-viral clearance.

Although CD8^+^ T cells are indispensable for influenza control, the role of CD8^+^ T cells in lung pathology is increasingly reported (*9, 13–15*). Data from the 2009 H1N1 pandemic show a strong correlation of the increased numbers/responses of CD8^+^ T cells with influenza disease severity (*16*). However, mice with impaired CD8^+^ T cell responses eventually succumb to the infection due to their inability to control the viral load (*17*). Thus, paradoxically, CD8^+^ T cells while indispensable to influenza control, also contribute to the significant lung pathology that exacerbates the disease. These responses rely on the development of cytotoxic CD8^+^ T cells armed in cytotoxic molecules such as perforin, and granzyme B (GzmB), that exert a cytolytic (CTL) function to kill viral-infected cells (*18–20*). However, the magnitude of CD8^+^ T cell cytotoxic function needs to be appropriately gaged for an effective virus removal without causing excessive collateral damage (*13*). A recent report of the influenza-specific cytotoxic CD8^+^ T cells, causing bystander damage to non-infected alveolar epithelial cells (*14*), provides evidence that the balance between protective and pathological cytotoxic functions can be easily disturbed. Besides their direct interaction with epithelial cells, CD8^+^ T cells produce a range of inflammatory cytokines that can promote inflammation and damage structural cells, compromising the lung barrier (*14, 21, 22*).

Interferons (IFNs) are crucial to regulating CD8^+^ T cell responses *via* interferon-inducible chemokines, C-X-C motif chemokine 9 (CXCL9) and 10 (CXCL10) (*23*), and consequently, regulate the CXCR3 dependent effector function (*24, 25*). While the role of CXCR3 has been described in the context of influenza CD8^+^ T cell effector/memory generation, there is a significant knowledge gap if CXCR3-CD8^+^ T cell axis regulates the development of pathologic host responses during primary influenza infection. Additionally, it remains to be determined if IFN types differentially regulate the development of CD8^+^ T cytotoxic responses during primary influenza infection. Therefore, the pathologic function of CD8^+^ T cells in the context of influenza disease warrants further investigation.

In this study, we sought to elucidate the role of CD8^+^ T cells during influenza infection, using a mouse model of severe but non-lethal influenza that displays a robust lung injury during the peak viral load at 7 days post-infection (dpi) as well as in post-viral resolution period (14 dpi), mimicking severe human influenza with persisting lung injury. We characterized the CD8^+^ T cell transcriptional diversity and studied the regulation of CD8^+^ T cell pathways in the lung during the peak viral load and viral resolved phases. We found that CD8^+^ T cells expressing IFN-inducible chemokine receptor, CXCR3 (Cxcr3^hi^ CD8^+^ T cells), exhibited a dominant enrichment of IFN-I induced molecular pathways, produced higher levels of cytolytic molecules, and persisted in the lung even after viral clearance at 14 dpi. The antibody-mediated neutralization of CXCR3 resulted in reduced lung injury, disease severity, and faster resolution of injury in virus-cleared lungs. These findings provide strong support that the CXCR3 pathway is dispensable for influenza clearance but mediates influenza-associated lung pathology and suggest that interference with this pathway could be explored therapeutically.

## RESULTS

### Activated CD8^+^ T cells persist in the lung after the viral clearance and correlate with lung injury

To establish spatiotemporal relation between lung pathology and the presence of CD8^+^ T cells, we conducted studies in the severe influenza model, characterized by significant lung inflammation, airway, and vascular damage with a substantial loss of body weight (∼ 25%). The inflammation and acute lung damage were determined by a combined metric of H&E staining of lung sections and the levels of lactate dehydrogenase (LDH) and albumin in the bronchoalveolar lavage (BAL) fluid. While the measure of acute lung injury, LDH, and albumin levels peaked at 7 dpi and normalized by 14 dpi, the histological analysis showed sustained lung inflammation and consolidation throughout 14 dpi before substantial resolution by 21 dpi (fig. S1, A to G). Influenza viral load in the lung was cleared by day 10 (fig. S1H). We next characterized the CD8^+^ T cells in mock and influenza-infected lungs during different stages of lung injury, at 3-21 dpi. The CD8^+^ T cells demonstrated a peak quantitative increase by 7 dpi, with their level remaining high until 14 dpi (Fig. 1A). Further analysis revealed that CD8^+^ T cells remained activated until 14 dpi (CD44^hi^ CD62^low^) (Fig. 1, B to D). Next, we characterized the effector and cytolytic function of CD8^+^ T cells by examining the intracellular expression of IFN-γ, perforin, and GzmB during different stages of influenza infection. The lung single cells from mock and influenza-infected mice were stimulated *in vitro* with influenza peptide NP_366-374_ (10μg/ml) for 6 hours before intracellular staining (*26*). We observed a peak of IFN-γ response on 7 dpi and its complete contraction by 14 dpi (Fig. 1, E and F). The expression of cytolytic molecules, perforin, and GzmB also peaked at 7 dpi. However, contrary to the IFN-γ positive CD8^+^ T cells, perforin and GzmB expressing cells were detected in the lung well beyond the viral clearance timepoint (>day 14) (Fig. 1, G to J).

**Fig. 1.**
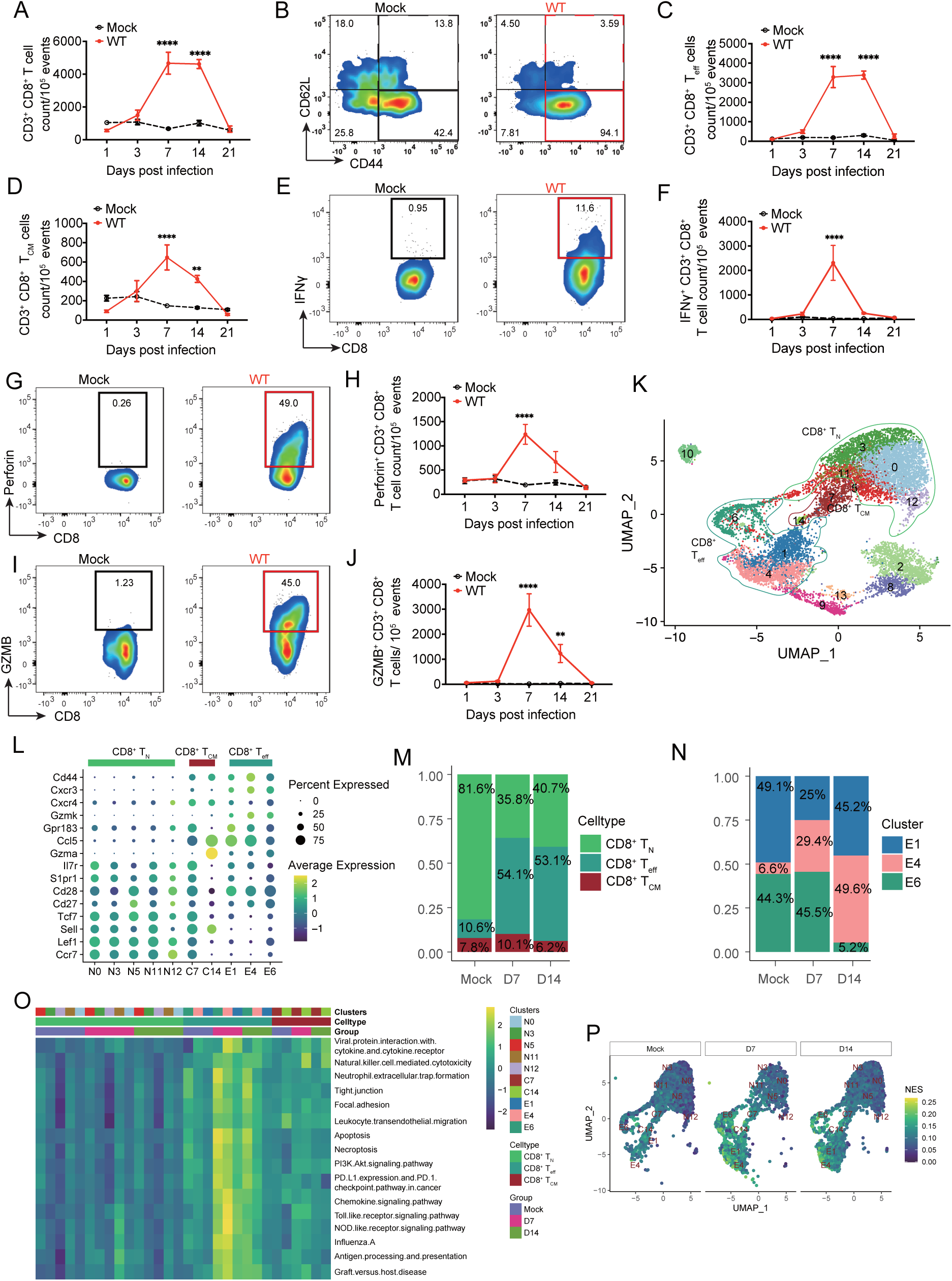
Cellular diversity and relative distribution of lung CD8^+^ T cells in influenza model. WT (C57BL/6) mice were mock-infected with PBS or infected with 1,000 PFUs of IAV (PR8) intranasally. At indicated time points, mice were euthanized, the BAL fluid and lungs were aseptically collected. The FACS sorted CD8^+^ T cells from mock and IAV-infected mice were subjected to scRNAseq analysis and lung single cells were used for flow cytometry characterization of CD8^+^ T cells. **(A)** Kinetics of CD8^+^ T cells. **(B)** Representative graphs showing the gating for effector CD8^+^ T_eff_ cells (CD44^+^CD62L^-^) and central memory CD8^+^ T_CM_ (CD44^+^CD62L^-^) in mock (black boxes) and PR8-infected WT (red boxes) mice. **(C)** Count of CD8^+^ T_eff_ cells (CD44^+^CD62L^-^). **(D)** CD8^+^ T_CM_ (CD44^+^CD62L^-^) cells (D) from mock and PR8-infected (PR8) mice, with gating shown in (B). **(E-G)** FACS representative graphs and Kinetics of IFN-γ^+^ **(E** and **F),** Perforin^+^ **(G** and **H)** and Granzyme B^+^ (GzmB) (I&J)-expressing CD8^+^ T cells. Data pooled from 2 independent experiments in (**A** to **J)**: with n=10 per group per time-point. The data is shown as means ± SEM. Two-way ANOVA mixed comparison of the means was used for statistical significance, only significant differences at indicated time points are shown, * p< 0.05, ** p< 0.01, *** p< 0.005 and **** p<0 .001. **(K** to **P)** scRNAseq analysis of FACS sorted CD8^+^ T cells from mock and PR8-infected mice lung at 7 and 14 dpi. **(K)** Unbiased Uniform manifold approximation and projections (UMAP) of total CD8^+^ T cells from Mock and PR8-infected at 7 and 14 dpi, revealing 15 clusters of CD8^+^ T cells. (**L)** Dot plot representing expression levels of canonical markers genes for selected 10 clusters generated by unbiased analysis, and their corresponding subsets clustering within CD8^+^ T_N_, T_CM_ or T_eff_. Clusters were renamed as N for naïve, C for central and E for effectors corresponding with their CD8^+^ T cell subsets. **(M** to **N)** Distribution of the proportions of CD8^+^ T_N_, T_CM_ and T_eff_ clusters **(M)** and clusters E1, E4 and E6 **(N)**, corresponding to the CD8^+^ T_eff_ subset; and their kinetics at day 0 (Mock), 7 (D7) and 14 (D14) dpi. **(O)** Heat map of GSVA enrichment scores of selected significant pathways between CD8^+^ T_eff_ against T_N_ and T_CM_. Color stands for the up-regulated (yellow) or down-regulated (dark green) in cells. **(P)** GSVA enrichment scores of CD8^+^ T cells with UMAP embedding Natural killer cell mediated cytotoxicity pathway. Color stands for the normalized enrichment score (NES).

Since CD8^+^ T cells exhibited a persistent cytotoxic phenotype even in the virus-cleared lungs, we performed scRNA-Seq to study the cellular heterogeneity and regulation of cellular response in CD8^+^ T cells during peak viral (day 7) and post-viral resolution phase (day 14). After stringent filtration, the transcriptome data of a total of 12,406 high-quality cells (mock: 4,402, day 7: 3,553, day 14: 4,451) were retained for further analysis. Through unsupervised clustering, a total of 15 cell clusters were identified and visualized as uniform manifold approximation and projection (UMAP) embeddings (Fig. 1K). Five clusters (cluster 2, 8, 9, 10, and 13) were removed because of the low expression of *Cd8a* and *Cd8b1* (Fig. 1K and fig. S1I). Based on the canonical lineage-defining markers, ten cellular clusters were further grouped into three cell subsets as CD8^+^ T naïve (CD8^+^ T_N_: *Ccr7*, *Lef1*, *Sell*, *Tcf7*, *Cd27*, *Cd28*, and *S1pr1*), CD8^+^ T central memory (CD8^+^ T_CM_: *Il7r*, *Gmza,* and *Ccl5*) and CD8^+^ T effector/effector memory (CD8^+^ T_eff_: *Gzmk*, *Cxcr4*, *Cxcr3*, and *Cd44*) (Fig. 1, K and L, and fig. S1J) (*27*). We then compared the proportion of each cell subset during different stages of infection (7 dpi vs. 14 dpi) (Fig. 1M). In both mock and infection groups, T_eff_ subset fell into three cellular clusters, designated as E1, E4, and E6. Compared to the mock group, influenza infection led to a significant enrichment of T_eff_ cell clusters (E1, E4, and E6) at 7 and 14 dpi, whereas a simultaneous contraction of naïve subset (T_N_) was observed (Fig. 1M). To determine the relative ratio of T_eff_ cell clusters during different infection stages, we compared the cell proportion of the clusters E1, E4, and E6 in influenza-infected lungs at 7 and 14 dpi (Fig. 1N). Of these three clusters, cluster E6 had a similar abundance between Mock (44.3%) and 7 dpi (45.5%), and a dramatic reduction at 14 dpi (5.2%) (Fig. 1N). A significant contraction of cluster E1 was observed in influenza-infected lungs at 7 dpi (25%), and its restoration to almost the same level as mock by 14 dpi (45.2%) (Fig. 1N). The cluster E4 was significantly enriched at 7 dpi (29.4%, compared to mock-infection with 6.6%), and further expanded to become a dominant cluster at 14 dpi (49.6%) (Fig. 1N). These findings reveal that T_eff_ exhibits significant cellular diversity with dynamic changes in cellular clusters over time during influenza infection.

Next, we performed Gene Set Variation Analysis (GSVA) analyses to identify functionally enriched pathways in CD8^+^ T cells between mock and influenza-infected groups. Compared to CD8^+^ T_N_ and CD8^+^ T_CM_, pathways related to inflammation, cytotoxicity, and cell death were upregulated in CD8^+^ T_eff_ (Fig. 1O and data S1), at both at 7 and 14 dpi. Among those pathways, the “natural killer cell-mediated cytotoxicity” pathway was highly enriched in CD8^+^ T_eff_ (E1, E4, and E6) at 7 and 14 dpi after IAV infection (Fig. 1P). The scRNAseq findings, while concurring with flow cytometry data that cytotoxic CD8^+^ T cells persist in viral resolved lungs, provide another layer of evidence *vis-a-vis* the distribution of cellular heterogeneity and molecular response of CD8^+^ T_eff_ subsets. Because of our interest in understanding the CD8^+^ T cell pathologic response, we focused on T_eff_ cells as a subject of the in-depth scRNAseq analysis.

### Cxcr3 expression is linked with enhanced CD8^+^ T cell-specific cytolytic gene expression in influenza-infected lungs

The scRNAseq data show that the high expression of chemokine receptor *Cxcr3* (Cxcr3^hi^) was predominantly associated with T_eff_ subsets, in contrast with its low-level expression by T_CM_ and T_N_ CD8^+^ T cell subsets, designated as Cxcr3^low^ (Fig. 2, A and B). Furthermore, all-lung-leukocyte scRNAseq analysis revealed that CD8^+^ T cells represented the only cell type that highly expressed *Cxcr3* at 7 dpi (Fig. 2C). To examine the shared and unique differentially expressed genes (DEGs) between Cxcr3^hi^ and Cxcr3^low^ CD8^+^ T cells, we performed differential gene expression analyses from the mock and influenza-infected lungs at 7 and 14 dpi. A total of 614 DEGs between Cxcr3^hi^ and Cxcr3^low^ CD8^+^ T cells were common at these three sets (Fig. 2D and data S2). Notably, there were 109, 658, and 380 DEGs that were identified as unique DEGs (Cxcr3^hi^ vs Cxcr3^low^) at mock, 7, and 14 dpi, respectively. We also found 60 shared pathways that were significantly enriched among the DEGs, all of which were highly upregulated in Cxcr3^hi^ clusters in mock at 7 and 14 dpi (fig. S2A and data S3). Several pathways were linked to cytotoxicity and to infections, which are known to induce profound inflammatory damage.

**Fig. 2.**
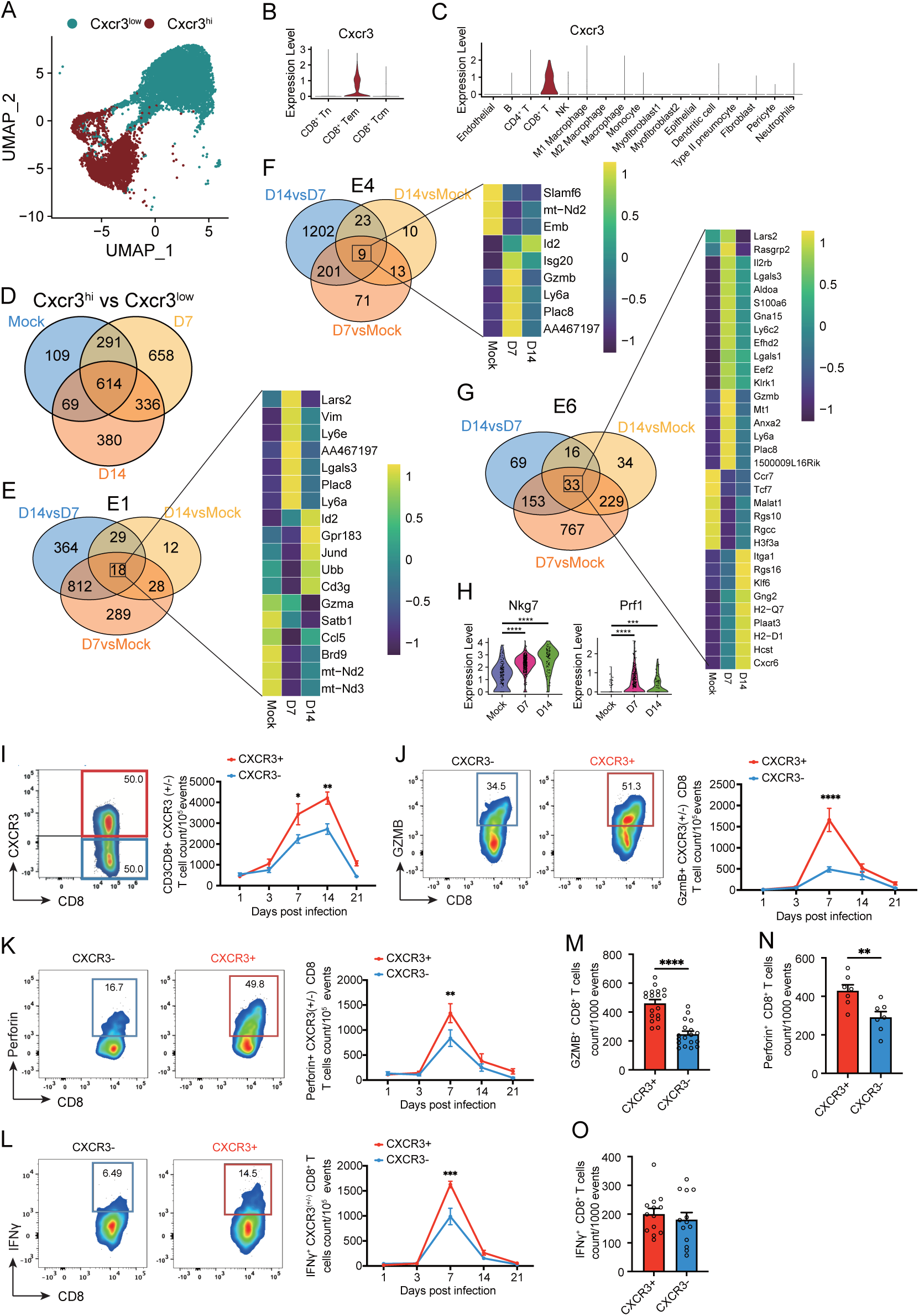
CXCR3 expression and CD8^+^ T cell response in influenza-infected lungs. WT (C57BL/6) mice were mock-infected with PBS or infected with 1000 PFUs of IAV (PR8) intranasally. At indicated time points, mice were euthanized, the BAL fluid and lungs were aseptically collected. The FACS sorted CD8^+^ T cells were subjected to scRNAseq analysis and lung single cells were used for flow cytometry characterization of CD8^+^ T cells. **(A)** UMAP embedding CD8^+^ single-cell transcriptomes from mock and PR8-infected mice annotated by *Cxcr3* expression. **(B)** Violin plots showing the expression *Cxcr3* within CD8^+^ T_N_, CD8^+^ T_eff_ and T_CM_ cell clusters. **(C)** The expression of *Cxcr3* was determined from a data set of scRNAseq from total lung cells of mock and PR8-infected mice at 7 dpi. **(D)** Venn diagram showing the distribution/number of shared and exclusively differentially expressed genes between Cxcr3^hi^ with Cxcr3^low^ CD8^+^ T cells from mock and PR8-infected mice at 7 and 14 dpi. **(E)** Venn diagram showing the numbers of the shared or unique differentially expressed genes identified between Mock vs D7 PR8, Mock vs D14 PR8, and D7 PR8 vs D14 PR8 in E1 cluster, and heatmap of the expression of 18 genes shared by the three comparisons within E1 cluster. **(F)** Venn diagram showing the numbers of the shared or unique differentially expressed genes identified between Mock vs D7 PR8, Mock vs D14 PR8, and D7 PR8 vs D14 PR8 in E4 cluster, and heatmap of the expression of 9 genes shared by the three comparisons within E4 cluster. **(G)** Venn diagram showing the numbers of the shared or unique differentially expressed genes identified between Mock vs D7 PR8, Mock vs D14 PR8, and D7 PR8 vs D14 PR8 in E6 cluster, and heatmap of the expression of 33 genes shared by the three comparisons within E6 cluster. Color bar indicates the expression level for each gene as indicated in each Venn diagram. **(H)** Violin plots showing the expression of *Nkg7* and *Prf1* in E6 cluster between Mock and PR8 at 7 and 14 dpi. Statistical significances were determined by MAST: *** p< 0.005 and **** p<0 .001. **(I** to **L)** FACS representative graph (Left) and Kinetics of CXCR3^+^ versus CXCR3^-^ CD8^+^ T cells **(I)**; representative graphs (left) and Kinetics (right) of GzmB**(J)**-, perforin**(K)**- and IFN-γ**(L)**-expression in CXCR3^+^ vs. CXCR3^-^ CD8^+^ T cells in PR8-infected WT mice at indicated time points. Data shown as means ± SEM, pooled data of 2 different experiments with n=10 per group per time point. Two-way ANOVA with mixed comparison of the means was used for statistical significance. **(M** to **O)** Number of CXCR3^+^ and CXCR3^-^ CD8^+^ T cells from PR8-infected mice were normalized at 1,000 events at 7 dpi. Expression of GzmB **(M)**, Perforin **(N)**, and IFN-γ **(O)** within CXCR3^+^ versus CXCR3^-^ CD8^+^ T cells were quantified. The data are representative of 2 independent experiments as in **M** to **O** and expressed as means ± SEM with n=10. Two-tailed T-test was used to compare the mean between CXCR3^+^ and CXCR3^-^ cells. For statistical significance: * p< 0.05, ** p< 0.01, *** p< 0.005 and **** p<0 .001.

Since Cxcr3^hi^ CD8^+^ T cells were associated with a dominant effector and cytotoxic response, we next investigated the cluster-specific host response in Cxcr3^hi^ CD8^+^ T cells. Venn diagrams represented the common and unique DEGs in Cxcr3^hi^ cell clusters (clusters E1, E4, and E6) in mock and influenza-infected lungs (Fig. 2, E, F and G). Eighteen shared DEGs were identified in cluster E1 among all three comparisons (Fig. 2E and data S4). Interestingly, *Gzma*, and *Ccl5* were highly expressed in mock Cxcr3^hi^ cluster E1 and a significant downregulation was observed in influenza-specific Cxcr3^hi^ cluster E1 at 7 and 14 dpi (Fig. 2E), whereas the expression of *Ly6a*, *Vim,* and *Ly6e* was highly expressed at 7 dpi, compared to mock or 14 dpi Cxcr3^hi^ cluster E1 (Fig. 2E). Gene Set Enrichment Analysis (GSEA) identified five significant pathways in all three comparisons. These include Pertussis, IL-17 signaling pathway, Hematopoietic cell lineage, B cell receptor signaling pathway, and Antigen processing and presentation (fig. S2B and data S5). Compared to mock, all five pathways were found to be upregulated at 7 and 14 dpi. The cluster E4 had 9 common DEGs between mock and 7 and 14 dpi infection time points (Fig. 2F and data S6). Among 9 DEGs, *Gzmb*, *Ly6a,* and *Plac8* were induced after influenza infection at 7 dpi, and a significant downregulation was observed at 14 dpi (Fig. 2F). A total of 22 shared pathways were significantly enriched in cluster E4 among all three comparisons (fig. S2C and data S7). Compared to clusters E1 and E4, cluster E6 had the highest number of DEGs, with 33 significant DEGs between mock and influenza-specific Cxcr3^hi^ cluster E6 at 7 and 14 dpi (Fig. 2G and data S8). Compared to mock and 14 dpi, cluster E6 exhibited higher expression of *Il2rb*, *Lgals3*, *S100a6*, *Lt6c2*, *Gzmb*, *Ly6a*, and *Plac8*. Significant downregulation of *Ccr7*, *Tcf7*, *Malat1*, *Rgs10*, *Rgcc*, and *H3f3a* was observed in influenza-infected lungs compared to mock at 7 and 14 dpi. We also found that cytotoxicity molecules *Nkg7* and *Prf1* were significantly upregulated at 7 and 14 dpi when compared to mock in cluster E6 (Fig. 2H). Furthermore, a total of 25 significant pathways were highly enriched in cluster E6 (fig. S2D and data S9), including those related to cytokine-cytokine receptor interaction, chemokine signaling pathway, and apoptosis. Overall, our data show the cellular heterogeneity in *Cxcr3* expressing CD8^+^ T cells, and cluster E6 being more profoundly associated with the expression of cytotoxic genes and pathways. We further validated the scRNAseq data to determine the functional differences between CXCR3^+^ and CXCR3^-^ CD8^+^ T cells by flow cytometry (Fig. 2, I to O). Consistent with these findings, CXCR3^+^ cells expressed significantly higher levels of GzmB and perforin, based on both total (Fig. 2, J and K) and normalized counts (per 1000 cells of CXCR3^+^ and CXCR3^-^) (Fig. 2, M and N). The CXCR3^+^ CD8^+^ T cells exhibited a peak cytotoxic response at 7 dpi and downregulation of response by 14 dpi (Fig. 2, M and N). Interestingly, apart from 7 dpi (Fig. 2L) both CXCR3^+^ and CXCR3^-^ CD8^+^ T cells expressed comparable levels of IFN-γ (Fig. 2, L to O), suggesting the central differences between CXCR3^+^ and CXCR3^-^ CD8^+^ T cells were related to their cytotoxic, rather than the effector functions.

### Unique molecular pathways and regulons associated with cytotoxic response in CXCR3^hi^ CD8^+^ T cells are revealed by scRNA-Seq analysis

To further unravel the differences between Cxcr3^hi^ and Cxcr3^low^ CD8^+^ T cell functional properties, we analyzed ligand-receptor interactions in the scRNAseq dataset. The ligand (outgoing signal) and receptor (incoming signal) in Cxcr3^hi^ and Cxcr3^low^ cellular clusters were compared among three groups, mock and influenza infection at 7 and 14 dpi. Compared to mock, CXCR3^hi^ cellular clusters (E1, E4, and E6) exhibited significant differences in incoming and outgoing signals at 7 and 14 dpi. Among the three CXCR3^hi^ clusters, the E4 cluster was the major source of incoming and outgoing signals at 7 and 14 dpi (Fig. 3A). Consistent with their enhanced cytotoxic properties, the CXCR3^hi^ clusters also exhibited higher ligand-receptor interactions at 7 and 14 dpi leading to the regulation of crucial signaling pathways in these clusters (Fig. 3, B and C). We identified 38 signaling pathways with significant ligand-receptor pairs in Cxcr3^hi^ and Cxcr3^low^ clusters, including THY1, PDL2, PD-L1, CLEC, CD86, CD6, CD39, ALCAM, PARS, IFN-I, MIF, CXCL, CCL, and SEMA4 (Fig. 3, B and C). To compare the patterns of outgoing (ligands) and incoming (receptors) signaling between Cxcr3^hi^ and Cxcr3^low^ clusters (mock, 7 and 14 dpi), we combined all identified signaling pathways from different datasets. We subsequently compared them in parallel, which allowed us to identify ligand-receptor pairing that exhibited different signaling patterns (Fig. 3, B and C). Compared with mock, the majority of pathways such as CCL, CXCL, MIF, IFN-I, PARs, ALCAM, CD39, CD6, CD86, CLEC, PD−L1, PDL2, and THY1 were found to be active in Cxcr3^hi^ cellular clusters at 7 dpi. Among the highly active pathways at 7 dpi, MIF, IFN-I, ALCAM, CD39, CD6, CLEC, and THY1 pathways were significantly downregulated by 14 dpi, except for cytotoxicity-triggering receptor NKG2D, which was upregulated in cluster E6 at day 14 pi (Fig. 3, B and C). In contrast, Cxcr3^low^ clusters exhibited enhanced IL-2 signaling (compared to Cxcr3^hi^ clusters) at 7 and 14 dpi (Fig. 3, B and C). Thus, Cxcr3^hi^ CD8^+^ T cells display unique molecular pathways associated with cytotoxic response in the lungs during influenza infection.

**Fig. 3.**
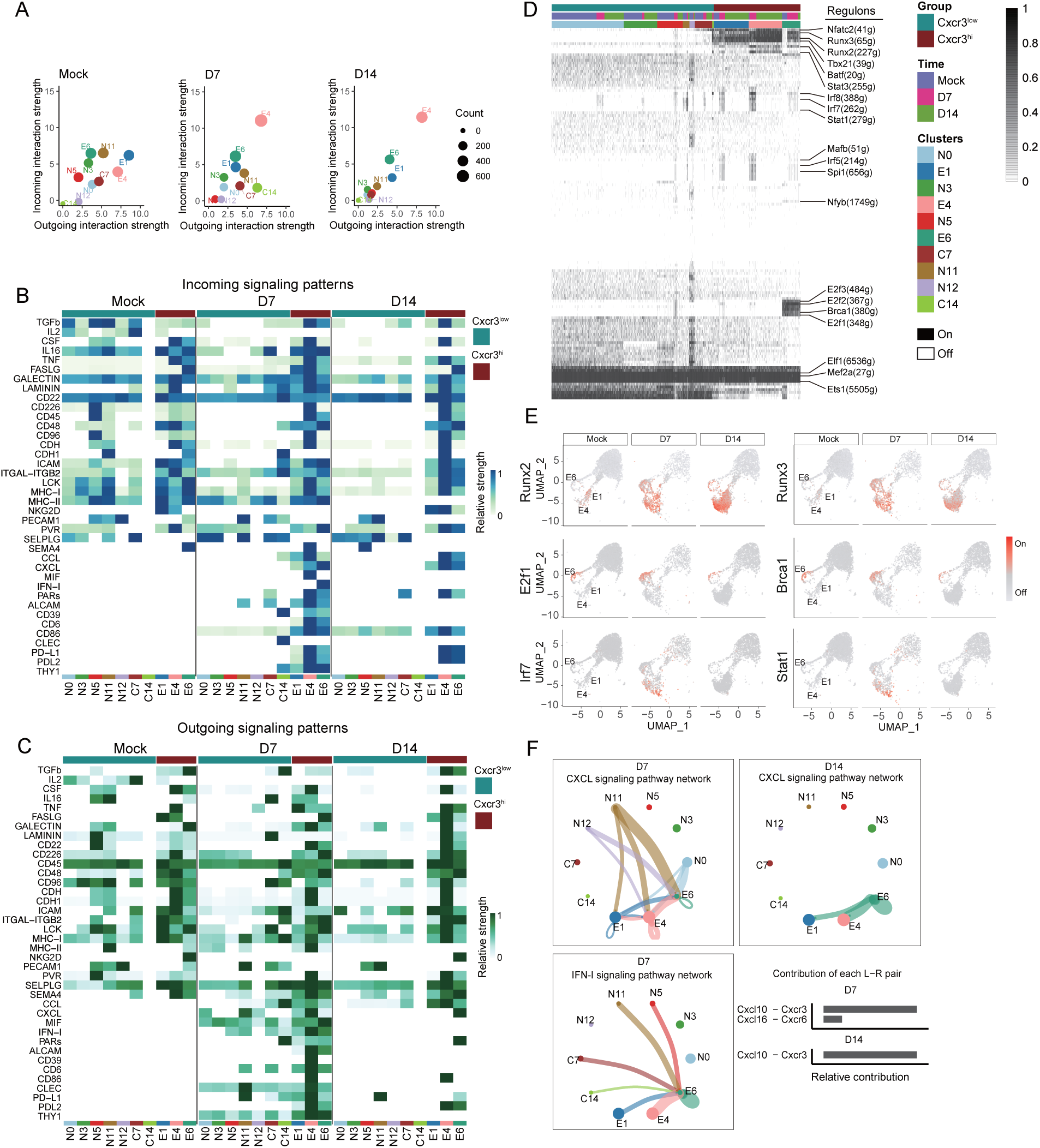
scRNAseq analysis of host response regulation in CXCR3^+^ CD8^+^ T cells. WT (C57BL/6) mice were mock-infected with PBS or infected with 1000 PFUs of IAV (PR8) intranasally. At indicated time points, mice were euthanized, the BAL fluid and lungs were aseptically collected. The FACS sorted CD8^+^ T cells were subjected to scRNAseq analysis. **(A)** Scatter plot showing the major source and targets in mock and PR8-infected mice at 7 and 14 dpi. The color indicates the cell types. Dot size is proportional to the number of inferred links (both outgoing and incoming) associated with each cell group. **(B** to **C)** Compare incoming **(B)** and outgoing **(C)** signaling associated with each CD8^+^ T cell cluster at 7 and 14 dpi. The colored bars show the relative importance of each cell group based on outgoing and incoming signaling patterns for pathways. **(D)** A heatmap of CD8^+^ cells showing high-confidence regulons at 7 and 14 dpi. mice. “On” indicates active regulons; “Off” indicates inactive regulons. Active regulons per cell appear in black; the horizontal color bar indicates the subset associated with each cell and group. Numbers in parentheses represent the number of genes comprising the regulon. **(E)** UMAP plots showing the activity of the indicated regulons. Cells in which the indicated regulon is active (regulon activity exceeds a regulon-specific area under the curve (AUC) threshold) are shown in red. **(F)** The inferred CXCL and IFN-I signaling networks and relative contribution of each ligand-receptor pair to the overall CXCL signaling network, respectively. Circle sizes are proportional to the number of cells in each cell group and edge width represents the communication probability.

Because cytotoxic CD8^+^ T cell response is regulated by a coordinated function of several transcriptional factors (*28, 29*), we examined the differences in transcriptional factors and their gene modules, also known as regulons, of Cxcr3^hi^ vs. Cxcr3^low^ CD8^+^ T cells using the single-cell regulatory network inference and clustering (SCENIC) software. A total of 227 regulons with 9,619 genes were identified across Cxcr3 clusters, which were further binarized and matched with cell types (Fig. 3D). Several important regulons, including IFN regulated *Tbx21*, *Nfatc2, and Batf,* were uniquely activated in Cxcr3^hi^ clusters. Additionally, *Runx3*, *Runx2*, and *Stat3* regulons were highly activated in Cxcr3^hi^ clusters (Fig. 3E), and significant downregulation of these regulons was observed in Cxcr3^low^ cells (Fig. 3E). Several regulons such as *Irf8*, *Irf7*, *Stat1*, *Mafb*, *Irf5,* and *Spi1* were commonly activated at 7 dpi and were turned off at 14 dpi in both Cxcr3^hi^ and Cxcr3^low^ cells (Fig. 3, D and E). Notably, consistent with the above findings of chemokine and IFN signaling, cluster E6 represented a majority of activated regulons in Cxcr3^hi^ clusters, suggesting that cluster E6 is a major contributor of inflammatory response in Cxcr3^hi^ CD8^+^ T cells (Fig. 3, D and E). Overall, these data identified several key regulons implicated in differential regulation of host response in Cxcr3^hi^ and Cxcr3^low^ clusters, further supporting that these cells are the major drivers of inflammation and cytotoxicity during influenza infection.

### CXCR3 pathway is a major element of the late-phase CD8^+^ T cell communication and source of IFN-I responsive CD8^+^ T cell subset recruited to the influenza lungs

Because chemokine signaling plays an instrumental role in shaping the CD8^+^ T cell responses (*30*), we determined the cellular communications for CXCL signaling in Cxcr3^hi^ and Cxcr3^low^ CD8^+^ T cells. Our data show that compared to mock, clusters N0, N11, and N12 from Cxcr3^low^ and clusters E1 and E4 from Cxcr3^hi^ cells were more profoundly associated with the expression of CXCL chemokine ligands at 7 dpi (Fig. 3F). However, E1, E4, and E6 were the main recipient clusters in Cxcr3^hi^ cells, while N0 was the only recipient cluster in Cxcr3^low^ cells at 7 dpi (Fig. 3F). The cluster E6 was the only expressor of CXCL chemokines at day 14 pi, and the clusters E1 and E4 acted as the recipient cells of Cxcr3^hi^ clusters at 14 dpi (Fig. 3F). We did not detect the chemokine signaling in Cxcr3^low^ cells at 14 dpi. The *Cxcr3*-*Cxcl10* and *Cxcr6-Cxcl16* ligand-receptor pairs were predominantly expressed at 7 dpi, suggesting that both pathways were involved in the early recruitment of CD6 T cells in the lungs. In contrast, the *Cxcl10*-*Cxcr3* ligand-receptor pair was the only contributor to the CXCL communication pathway at 14 dpi (Fig. 3F, bottom & right), suggesting that the late-phase CD8^+^ T cell recruitment was relying solely on the CXCR3 pathway.

We further examined the expression of CXCR3 ligand, *Cxcl9,* and *Cxcl10* in CD8^+^ T cell clusters. While *Cxcl10* expression was detected in each cluster (Cxcr3^hi^ and Cxcr3^low^), the expression of *Cxcl9* remained undetectable (fig. S3A), suggesting the CD8^+^ T cell-independent association of *Cxcl9* expression in our model. Next, we analyzed the ligand-receptor interactions of type-I (IFN-I) and type-II (IFN-II) interferons. While the clusters from both Cxcr3^low^ (N5, N7, N11, N14) and Cxcr3^hi^ (E1, E4, and E6) expressed IFN-I, the cluster E6 was the only cluster recipient of IFN-I signaling (Fig. 3F). We did not detect IFN-II receptor signaling among Cxcr3^hi^ or Cxcr3^low^ cell clusters in mock or IAV infected 7 and 14 dpi time points (fig.S3B). Thus, CXCR3/CXCL10 axis is the main pathway for both early and late CD8^+^ T-cell recruitment in the influenza-infected mice, and among Cxcr3^hi^ cells, cluster E6 appears to be the main driver of IFN-I dependent inflammatory response in Cxcr3^hi^ CD8^+^ T cells.

### The antibody blockade of CXCR3 mitigates influenza lung injury and disease severity without affecting viral clearance

Since CXCR3^+^ CD8^+^ T cells showed all the attributes of cells specialized in the most robust cytotoxic response, we postulated the CXCR3^+^ CD8^+^ T cells contributed to influenza lung injury. We performed antibody-mediated neutralization of CXCR3 in influenza-infected mice (Fig. 4A) and examined multiple parameters associated with lung inflammation, injury, and disease severity. The antibody-mediated neutralization of CXCR3 led to an approximately 70% reduction of overall CD8^+^ T cells in influenza-infected lungs at 7 dpi (Fig. 4, B and C). We did not observe a quantitative increase in the frequency of CD4^+^ T cells at 7 dpi, suggesting that CD8^+^ T cells acquired an early effector and cytolytic phenotype (Fig. 4, D and E) as early as 7 dpi. Moreover, CXCR3 blockade resulted in a significantly greater loss of CD8^+^ T cells than CD4^+^ T cells even at a late time point, 14 dpi. (Fig. 4E). These findings are consistent with our scRNAseq and flow cytometry data that CXCR3 is primarily expressed by CD8^+^ T cells in our model. The depletion of CXCR3 resulted in the reduced level of CCL2 chemokine and consequently the reduced CCR2 monocytes (Fig. 4, F and G). The administration of anti-CXCR3 antibody also resulted in significantly reduced levels of CD8^+^ T cell-specific effector (i.e., IFN-γ) and cytolytic molecules, (i.e., perforin, granzyme-B) (Fig. 4, H to J). Notably, while CXCR3 antibody blockade resulted in a significant loss of CD8^+^ T cell cytotoxic response, it did not abolish the CD8^+^ T effector and cytotoxic response altogether. We further measured the levels of CXCR3 cognate binding chemokines CXCL9 and CXL10, as well as cytokines in the lung homogenates of mock- and influenza-infected mice at 7 dpi. We found that CXCR3 antibody blockade resulted in reduced levels of CXCL9 and IL-10 (Fig. 4, K to L). The lower IL10 level is indicative of reduced lung inflammation in mice with CXCR3 antibody blockade. Overall, these data demonstrated that CXCR3 blockade dampened the expression of inflammatory and cytolytic molecules in influenza-infected lungs. To corroborate our CXCR3 antibody-based neutralization approach, we compared CD8^+^ T cell cytotoxic response in influenza-infected WT with those of CXCR3 deficient (CXCR3^-/-^) mice. The results demonstrated a significant loss of CD8^+^ T cells (approximately 70%) in the lung (fig. S4A) and a dramatic reduction in overall CD8^+^ T cell cytotoxic response (fig. S4, B to D) as well as decreased CCR2^+^ monocyte recruitment (fig. S4E) in CXCR3^-/-^ mice. Together, these data show that CXCR3 amplifies the magnitude of the inflammatory response during influenza pathogenesis, in particular, the recruitment and cytotoxic function of CD8^+^ T cells.

**Fig. 4.**
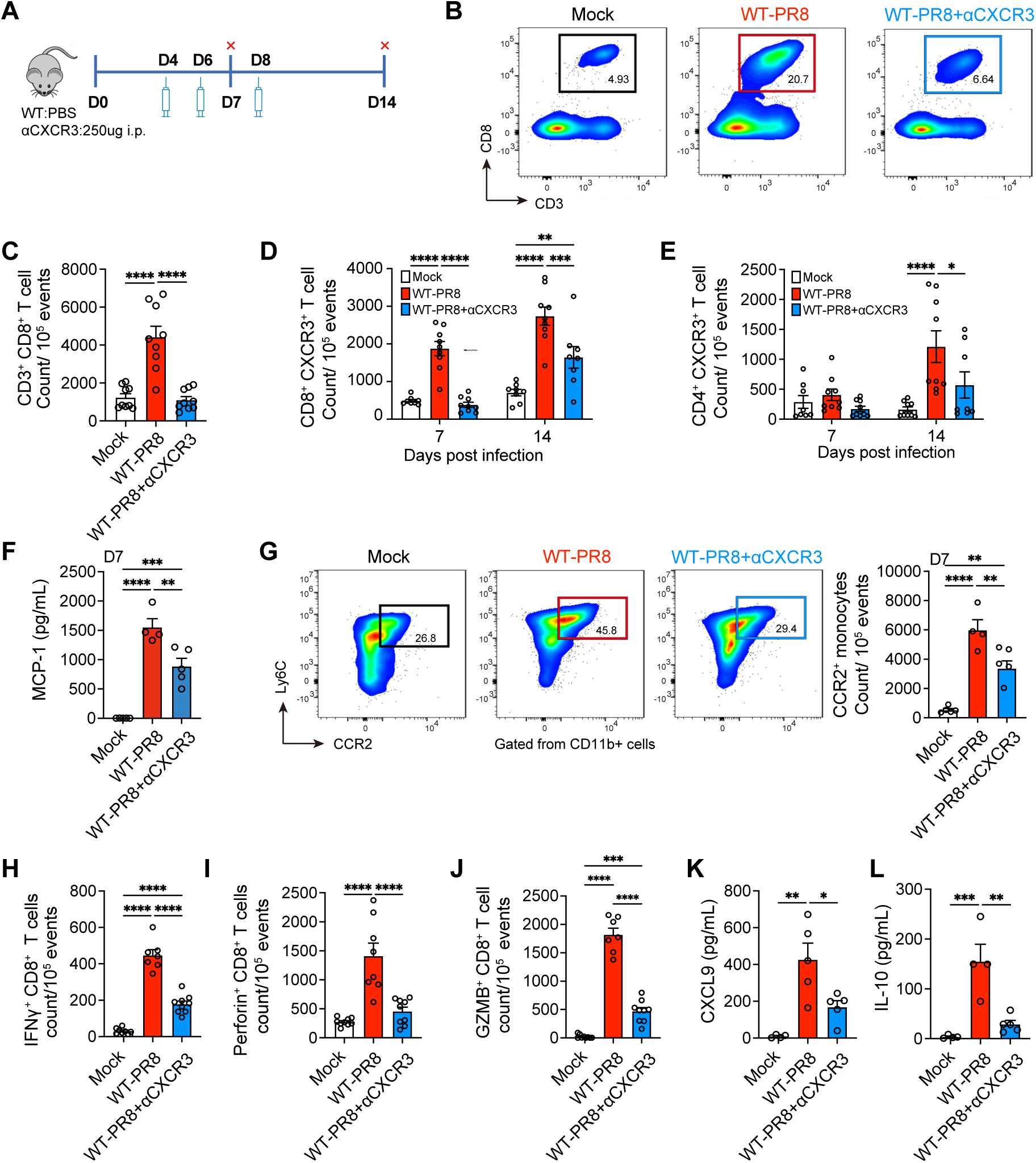
The effect of CXCR3 neutralization on CD8^+^ T cell cytotoxic responses and inflammation in IAV-infected lungs. **(A)** WT (C57BL/6) mice were mock-infected with PBS or infected with 1000 PFUs of IAV (WT-PR8) intranasally. PR8-infected mice received CXCR3 neutralizing antibody (WT-PR8+αCXCR3) every alternate day starting day 4 post-infection and mice were euthanized at 7 and 14 dpi. The levels of cytokines/chemokines were measured in homogenized lungs. The frequencies of immune cells were determined in lung single cells using flow cytometry. **(B** to **C)** Flow cytometry representative graphs **(B)** and **(C)** number/quantification **(C)** of CD8^+^ T cells at 7 dpi. **(D** to **E)** Count of CXCR3^+^ CD8^+^ T cells **(D)** and CXCR3^+^ CD4^+^ T cells **(E)** at 7 and 14 dpi. **(F)** Levels of MCP1/CCL-2 at 7 dpi, measured using BiolegendPlex kit. Data are shown as means ± SEM, representative data of 2 experiments with n=5 per group. **(G)** Flow cytometry representative graphs (left) and count of CCR2^+^ monocytes at 7 dpi. CCR2^+^ monocytes were gated from CD45^+^ CD11b^+^ Ly6C^+^ cells. **(H** to **J)** Count of IFN-γ **(H)**, Perforin **(I)** and Granzyme B (GzmB) **(J)**-expressing CD8^+^ T cells at 7 dpi. Data are shown as means ± SEM, pooled data of 2 experiments with n=8-10 per group. **(K** to **L)** Levels of CXCL9**(K)** and IL-10**(L)** at 7 dpi, measured using BiolegendPlex kit. Data shown as means ± SEM with n=5. One-way ANOVA with Tukey post hoc test for multiple comparison (means) was used for statistical significance in (**C** to **L)**. * p< 0.05, ** p< 0.01, *** p< 0.005 and **** p< 0.001.

We further hypothesized that CXCR3 depletion would improve/reduce inflammation and pathology while sustaining antiviral responses through the function of CXCR3^-^ CD8^+^ T cells. We compared multiple parameters of influenza disease severity, i.e., weight loss, inflammation, and acute lung damage, in influenza-infected mice treated with or without anti-CXCR3 antibody. Compared to influenza-infected control mice (WT-PR8), mice treated with CXCR3 antibody (WT-PR8-αCXCR3) exhibited significantly reduced lung injury, evidenced by a reduced level of LDH in BAL (Fig. 5A) and reduced inflammation shown in and H&E tissue pathology at 7 and 14 dpi (Fig. 5, B to D). CXCR3-neutralized infected mice also exhibited reduced vascular damage and lung permeability based on higher expression of platelet endothelial cell adhesion molecule (CD31) (immunofluorescence lung sections) (Fig. 5E) and reduced leakage of serum albumin level in the BAL (Fig. 5G). Furthermore, CXCR3 blockade ameliorated bronchial epithelial injury, evidenced by improved anti-EpCAM (epithelial) immunofluorescence staining of lung sections (Fig. 5F). While LDH and albumin levels peaked at 7 dpi and significantly reduced by 14 dpi, the H&E tissue-pathology assessments showed a non-resolving lung injury and vascular damage in influenza-infected mice even at 14 dpi. Consistent with reduced lung inflammation and injury, CXCR3-neutralized mice also exhibited a significantly reduced weight loss (Fig. 5H). These findings were subsequently validated in CXCR3^-/-^ mice showing similarly reduced lung inflammation and pathology evident on histological sections and through the reduced levels of LDH and albumin in the BAL compared to the WT mice (fig. S5, A to C). These data demonstrated that the CXCR3 pathway is an important driver of lung injury during the peak viral load (7 dpi) and its disruption facilitates the expedited resolution of lung injury following the viral clearance. Since abolished CXCR3 signaling had such a profound impact on limiting lung pathology, we assessed its role in viral clearance to explore its usefulness as a therapeutic target. Both anti-CXCR3 antibody neutralization (Fig. 5I) and CXCR3 gene deletion (fig. S4C) did not affect viral clearance throughout the studied course of infection. These data consistently showed that, while CXCR3-signaling is an important driver of the immune-mediated lung pathology, it is dispensable to the clearance of influenza virus.

**Fig. 5.**
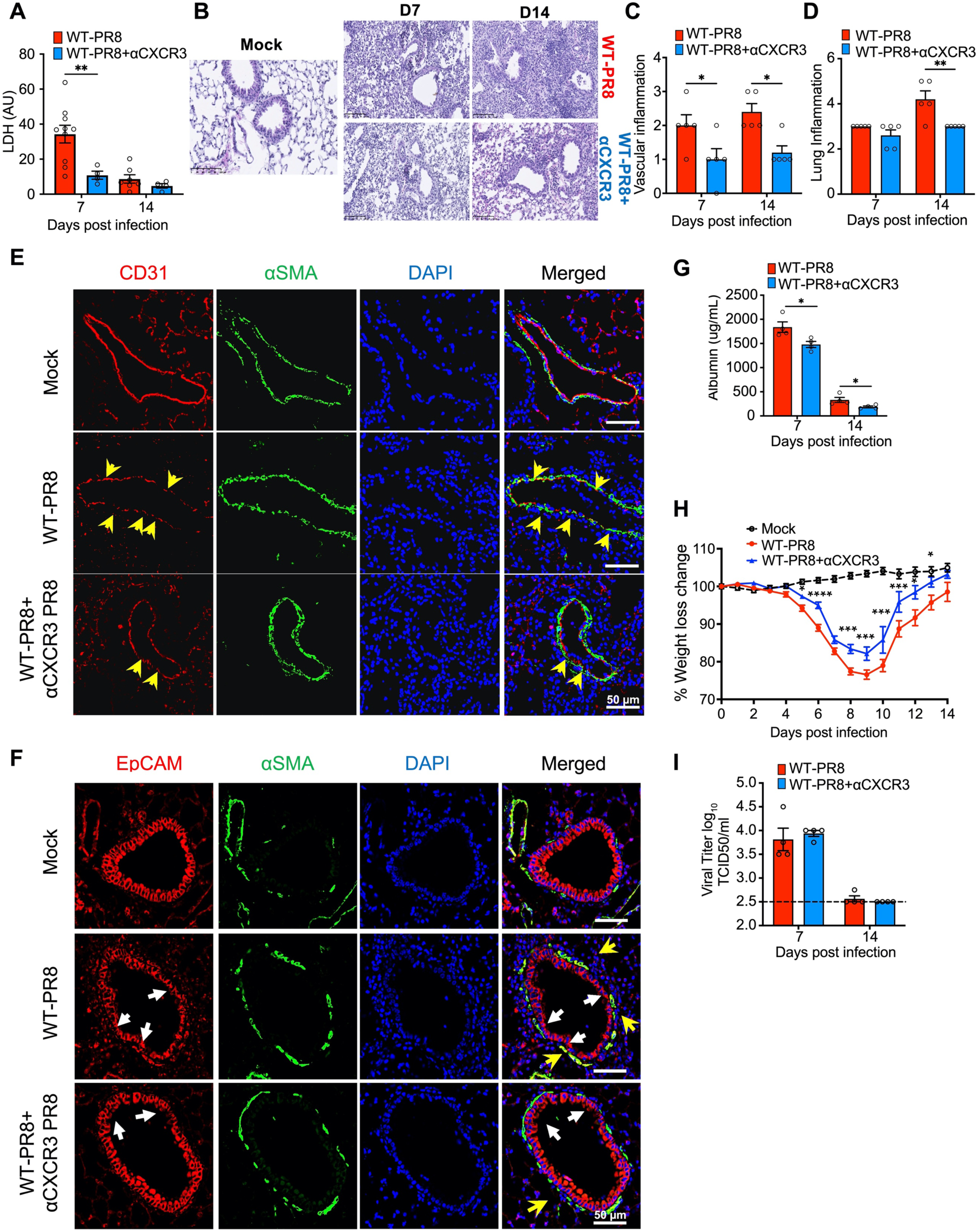
The effect of CXCR3 depletion on lung inflammation and pathology in influenza model. WT (C57BL/6) mice were mock-infected with PBS or infected with 1000 PFUs of IAV (WT-PR8) intranasally. PR8-infected mice received CXCR3 neutralizing antibody (WT-PR8+αCXCR3) every alternate day starting day 4 post-infection and mice were euthanized at 7 and 14 dpi. **(A)** LDH level in the BAL fluid at 7 and 14 dpi. Data is representative of 2 experiments, with n=5-10 mice per group/time point and shown as means ± SEM. Statistics were calculated using One-Way ANOVA with Tukey’s post hoc analysis. **(B)** Lung H&E staining at 7 and 14 dpi. Representative images of 2 independent experiments. n=5 per group/time point, scale bar 100 um. **(C** to **D)** Vascular/endothelial**(C)** and lung**(D)** inflammatory scores of histological analyses from H&E-stained lung sections. Data is representative of 2 independent experiments, n=5 per group/time point. Mann-Whitney test was used to compare the median between groups at 7 or 14 dpi. (**E** to **F)** Immunofluorescent staining of lung sections for endothelial/vascular **(E)** and epithelial **(F)** damage control at 7 dpi. Mock (top) and PR8 infected mice (WT-PR8) (middle) and PR8 infected mice recipient of αCXCR3 (WT-PR8+αCXCR3) (bottom). **(E)** CD31 (red) for endothelial cells, αSMA (green) alpha Smooth muscle cells; CD31 (red) to highlight the integrity of endothelial cell’s layer; where yellow arrowheads show the loss of endothelial cells marked by the discontinuity of red line following PR8-infection. EpCAM (red) was used in **(F)** to probe airway broncho-epithelial cells. White arrows highlight the loss of epithelial layer integrity, and yellow arrows the infiltrating inflammatory cells around the airways. **(G)** BAL albumin at 7 and 14 dpi. T-test was used at each time point. Data is representative of 2 independent experiments, with n=5 per group. Statistics were calculated using One-Way ANOVA with Tukey’s post hoc analysis. **(H)** Kinetics of body weight change at 1-14 dpi. Data are shown as pooled data of two independent experiments with n=10 per group. Statistics were calculated using Two-way ANOVA with Tukey post hoc test. **(I)** TCID_50_ Viral titer in lung homogenates at 7 and 14 dpi. Data are representative of two independent experiments with n=4 per group per time point. Black dashed line indicates the baseline value (mock) negative control. Statistics were calculated using One-Way ANOVA with Tukey’s post hoc analysis. * p< 0.05, ** p< 0.01, *** p< 0.005 and **** p< 0.001.

### IFN-γ is dispensable to the recruitment and regulation of cytotoxic response of CXCR3^+^ CD8^+^ T cells

The CXCL9 and CXCL10 chemokine gradients promote the recruitment of CXCR3^+^ T cells in the infected or inflamed tissues (*31, 32*). Influenza infection kinetics show that the levels of CXCL9 and CXCL10 peak in the lung at 7 dpi (Fig. 6, A and B). CXCL9 and CXCL10 are interferon-inducible chemokines (*33*), and we observed a significant correlation of CXCL9/10 protein level with those of IFN-γ in the BAL of influenza-infected mice (Fig. 6C). To further determine the cellular sources of CXCL9 and CXCL10 in our model, we analyzed the scRNAseq data of total lung cells from mock and influenza-infected mice. The scRNAseq data show that both hematopoietic and non-hematopoietic cells were significant expressors of *Cxcl9* and *Cxcl10* (Fig. 6D).

**Fig. 6.**
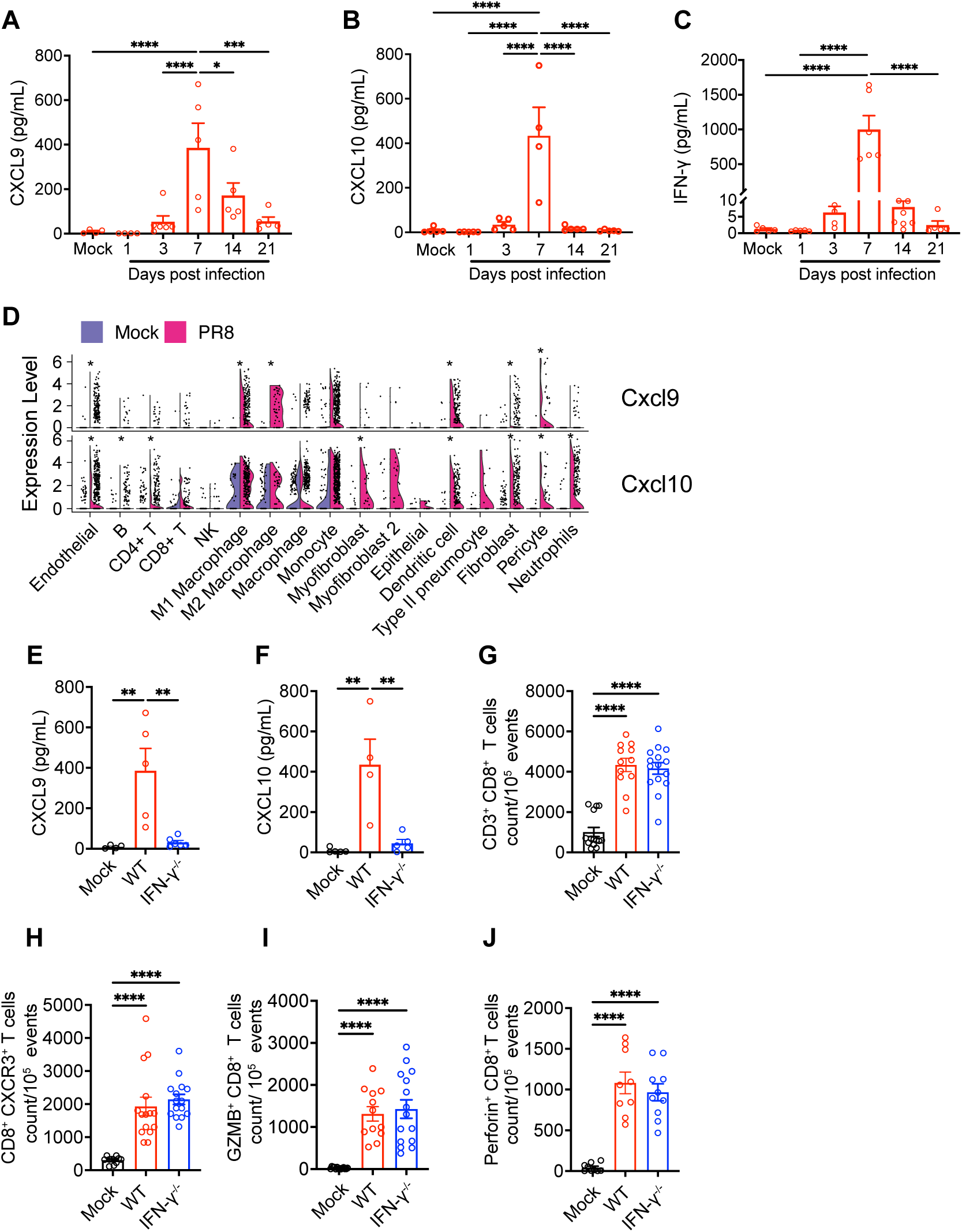
The role of IFN-γ in regulating the CXCR3^+^ CD8^+^ T cells response in influenza model. WT (C57BL/6) or Interferon-gamma deficient (IFN-γ^-/-^) mice were mock-infected with PBS or infected with 1000 PFUs of IAV (PR8) intranasally. Mice were euthanized at 7 and 14 dpi. The levels of cytokines/chemokines were measured in homogenized lungs. The frequencies of immune cells were determined in lung single cells using flow cytometry. **(A** to **C)** Kinetics of CXCL9 **(A)**, CXCL10 **(B)**, and IFN-γ **(C)** at indicated time points. The data is representative data of 2 independent experiments with n=5. **(D)** scRNAseq analysis of total lung single cells of mock and PR8-infected WT mice at 7 dpi. Violin plots showing the expression of CXCR3 ligands CXCL9 and CXCL10 in lung cells. *Mean significant difference between mock and IAV-infected mice. **(E-F)** Levels of CXCL9 **(E)** and CXCL10 **(F)** at 7 dpi. **(G** to **H)** Number of total **(G)** and CXCR3^+^ CD8^+^ T cells **(H)** at 7 dpi. **(I-J)** Expression of GzmB **(I)** and Perforin **(J)** in CD8^+^ T cells. **(G** to **J**) The data is representative of 2 independent experiments with n=10. One-way ANOVA with Turkey post hoc test was used for statistical significance. Data shown as means ± SEM, * p< 0.05, ** p< 0.01, *** p< 0.005 and **** p<0 .001.

We further addressed whether the IFN-γ signaling via the induction CXCL9/10 was responsible for the recruitment and enhanced cytotoxic response of CXCR3^+^ CD8^+^ T cells. Although IFN-γ deficiency led to a significant reduction in the levels of CXCL9 and CXCL10, it did not impair the recruitment of CD8^+^ T cells, as both WT and IFN-γ^-^/^-^ mice exhibited similar levels of CXCR3^+^ or total CD8^+^ T cell frequency in influenza-infected lungs (Fig. 6, E to H). These data suggest that IFN-γ-independent chemokines likely compensate for the lack of IFN-γ in recruiting CD8^+^ T cells. Our CD8^+^ T cell scRNAseq data show a dominant interferon signaling that correlated with exuberant cytotoxic response in Cxcr3^hi^ CD8^+^ T cells. We, therefore, investigated if IFN-γ regulated the cytotoxic function of CXCR3^+^ CD8^+^ T cells. We performed intracellular cytokine staining to investigate the cytolytic (GzmB, Perforin) properties in CD8^+^ T cells. The CD8^+^ T cells from WT and IFN-γ^-^/^-^ mice (influenza-infected) did not exhibit any difference in the intracellular expression of GzmB and Perforin (Fig. 6, I and J). These data suggest that IFN-γ is dispensable to the recruitment and regulation of cytotoxic molecules expression in CXCR3^+^ CD8^+^ T cells during influenza pathogenesis.

## DISCUSSION

In this study, we dissected the CD8^+^ T cells responses in influenza-infected lungs during the peak viral (acute) and virus-cleared states. The major finding of this study is that the CD8^+^ T-cell population recruited to the influenza-infected lungs represent significant transcriptional and functional diversity, with a subset not required for viral clearance but instead driving the severe lung pathology. The pathological CD8^+^ T-cell subset is characterized by high CXCR3 expression, the enhanced cytotoxic pathway signature, and their persistence in the lungs resulting in increased lung epithelium and vascular damage and the extended time of inflammatory infiltrate lung consolidation. Intercepting CXCR3 with either antibody or genetic deletion prevented the development of the severe influenza lung pathology, without affecting viral clearance. Thus, the temporal blocking of the CXCR3 pathway could be a viable candidate for therapeutic intervention that may prevent the development of significant lung injury during influenza-induced pneumonia.

While a protective role of CD8^+^ T cells against influenza remains unequivocally indispensable (*18, 26, 34, 35*), a growing body of literature suggests that exuberant CD8^+^ T cell responses might implicate lung injury and exacerbate disease severity (*9, 16*). The CD8^+^ T cells can cause damage to non-hematopoietic cells by direct interaction of cytotoxic cells with infected and uninfected cells (*36*). Sandt *et al.* showed influenza-specific human CD8^+^ T cells cause bystander damage to non-infected epithelial cells leading to the disruption of epithelial barrier integrity (*14*). Therefore, the goal of this study was to conduct a detailed analysis of CD8^+^ T cells to gain an insight into their role in both viral clearance and the development of influenza-associated lung pathology. To this end, we used a combined approach involving scRNAseq and flow cytometry to unravel the cellular diversity and regulation of inflammatory response in CD8^+^ T cells and their role in maintaining a balance between viral clearance and tissue pathology. We found that the CD8^+^ T cell functional heterogeneity can be distinguished based on the expression of the chemokine receptor, CXCR3. While Cxcr3^hi^ CD8^+^ T_eff_ cells are associated with enhanced cytotoxic response, CD8^+^ T with low *Cxcr3* expression (T_CM_), while positive for perforin and granzyme, show less pronounced expression of cytotoxic molecules and cytotoxic pathways. However, based on IFN-γ^+^ effector response, both Cxcr3^hi^ and Cxcr3^low^ CD8^+^ T cells displayed equally potent effector phenotype, suggesting that *Cxcr3* expression did not dictate the development of effector response in CD8^+^ T cells. Instead, the *Cxcr3* expression was a significant determinant of cytotoxic response in CD8^+^ T cells.

Little is known about the role of CXCR3 signaling in influenza. Fadel *et al.* showed that while CXCR3 deficiency protected the CCR5 deficient mice from influenza mortality, CXCR3 deficiency on its own did not affect the survival of influenza-infected mice in a lethal challenge model (*37*). In contrast, another study showed that CXCR3 deficient mice had increased survival following lethal influenza challenge, and neutrophils were reported to be the primary CXCR3 expressing cells (*38*). These contrasting data necessitate further evaluating the role of CXCR3 as a pathologic framework in influenza lung injury. Our model is different from those prior studies because we used a severe but non-lethal model that allowed us to study the lung injury during both the peak viral load and resolution. It is crucial because post-influenza complications involve persistent lung injury and impaired repair with compromised lung functions after the virus is already cleared (*39, 40*). A similar phenomenon of persistent lung injury following viral clearance is observed in ongoing coronavirus disease (COVID) pandemics (*41*). In this context, antibody blockade of CXCR3 ameliorated lung injury during the peak viral titers and led to a faster resolution of post-infection lung injury. We did not detect CXCR3^+^ neutrophils in our model. Our scRNAseq data from total lung cells demonstrated that CD8^+^ T cells were the only significant cell type associated with CXCR3 expression during the peak viral load in the lung.

Interferons are crucial in regulating the anti-viral function of CD8^+^ T cells (*42–44*). In particular, IFN-γ is a key regulator of the chemokines CXCL9 and CXCL10 that recruit CXCR3^+^ CD8^+^ T cells to the site of infection (*23*). In our model, despite being a crucial regulator of CXCL9 and CXCL10 chemokines, IFN-γ was found to be dispensable to the recruitment of CXCR3^+^ or total CD8^+^ T cells. Our data agree with prior reports showing that IFN-γ deficiency did not impact the recruitment of CD8^+^ T cells in influenza models (*45*), suggesting the IFN-γ-independent chemokine signaling in driving the CD8^+^ T cell recruitment in influenza-infected lungs. Furthermore, similar expression levels of perforin and granzyme-B in CD8^+^ T cells from WT and IFN-γ^-^/^-^ mice suggest the dispensability of IFN-γ in regulating the expression of cytolytic molecules in CXCR3^+^ CD8^+^ T cells. These data are further substantiated by the scRNAseq analysis of cell-cell communication, demonstrating the weak ligand-receptor interactions for IFN-II in CXCR3^hi^ CD8^+^ T cell clusters. Instead, the scRNAseq revealed enrichment of IFN-I receptor signaling and activation of multiple IFN-I-dependent regulons in Cxcr3^hi^ CD8^+^ T cells. Remarkably, our scRNAseq analysis identified the functional diversity in CXCR3^+^ CD8^+^ T cells with a specific subset E6 exhibiting a major contributor to host responses, including the enrichment of IFN-I signaling in CXCR3^+^ CD8^+^ T cells. These findings suggest the key role of IFN-I receptor signaling as a potential determinant of the inflammatory response of Cxcr3^hi^ CD8^+^ T cells that warrant further investigation.

In summary, this study unveiled the CD8^+^ T cellular heterogeneity associated with pathologic host response in influenza model during active infection with peak viral and during viral-cleared infection recovery. Our findings have shown that the CXCR3 blockade approach benefited the host improving inflammatory balance in the lung, preventing excessive host response-driven lung injury during the peak viral load and a faster resolution of lung injury without inhibiting viral clearance. These data suggest that CXCR3 pathway interception could be explored clinically in patients with severe influenza as a treatment option for reducing lung damage and accelerating post-influenza recovery.

## MATERIAL AND METHODS

### Mice and Influenza infection model

Wild-type (WT) C57BL/6J, CXCR3^-/-^, and IFN-γ^-/-^ mice were originally bought from the Jackson Laboratory and bred in-house. An equal proportion of 6–8-week-old sex-matched mice were included in this study. All experiments were performed according to the approved protocol by the University of North Dakota Animal Care and Use Committee (IACUC) (protocol #1808-8). Mice received food and water *ad libitum* at all times. Influenza A Virus (H1N1 A/Puerto Rico/8/1934 or PR8) was purchased from Charles River, Norwich, CT, and a plaque assay (*46*) was performed to determine the plaque forming units (PFUs) for IAV. For IAV infection, mice were lightly anesthetized with 4%v/v isoflurane/oxygen mixture and intranasally inoculated with 1000 PFUs of IAV in 50 µl volume. At indicated time points (days 3-21), mice were euthanized by CO_2_ exposure followed by cervical dislocation, and then lungs and BAL fluid were aseptically isolated and processed for downstream applications.

### Histopathology

Mice were euthanized and after perfusion, the left lobe was fixed in 10% neutral buffered (pH 7.4) formalin for 24 hr at room temperature prior to transferring into 70% ethanol. The lung tissues were embedded in paraffin, sliced into 5-mm sections to reveal the maximum longitudinal view of the main intrapulmonary bronchus of the left lobe, and stained with hematoxylin and eosin (H&E). H&E staining was performed by the histology core, University of North Dakota. Lung inflammation was evaluated and quantitatively by 2 blinded pathologists, on a scale of 0-5, with increments of 0.5 (15), with 0 as no inflammation and 4 as the highest degree of tissue infiltration of immune cells. Three parameters were focused in H&E analysis, Inflammation index (alveolar inflammation, inflammatory cell infiltration), bronchial epithelial damages (thickness, thinning, necrosis of bronchial epithelium, mucosal and inflammatory plugs), and vascular endothelium change (*47*).

### BAL Albumin and Lactate Dehydrogenase

Bronchial lavage fluid (BALF) was collected at each endpoint by instillation and aspiration of lung with 1ml of sterile ice-cold PBS through a G20 tracheal catheter (BD Biosciences). BALF supernatants were collected and stored -80-C until use. Albumin concentration in BAL samples was determined using Albumin ELISA kit (ALPCO, Salem, NH) following the manufacturer’s instructions. The level of lactate dehydrogenase (LDH) was measured in the BALF of mock and IAV-infected mice using a colorimetric LDH Assay kit following the manufacturer’s recommendations (ab102526, Abcam).

### Measurement of cytokines and chemokines

At indicated time points, lungs were homogenized in tissue protein extraction reagent using the tissue homogenizer 850 (Fischer scientific). The levels of cytokines and chemokines (CXCL9, CXCL10, MCP1) were determined in the lung lysates or BALF samples using LegendPlex mouse Proinflammatory chemokine panel (13-plex, Cat # 740451) and mouse inflammatory panel (13-plex, Cat # 740446) respectively, following the manufacturer’s instructions. Samples were acquired on a BD FACSymphony A3 flow cytometer, and data were analyzed using LEGENDplex V8.0 Data Analysis Software (BioLegend).

### Flow cytometry

The lungs were aseptically collected from mock and IAV-infected mice, and single cells were prepared after collagenase digestion of lungs, as previously described (PMC7336542). One million cells per sample were stained with Ghost Dye-BV510 (Tonbo Biosciences, San Diego, CA) and anti-mouse CD16/CD32 (FC blocked) (BD Biosciences, San Jose, CA) for 30 min at 4°C for 30 min. For cell surface staining, cells were incubated for 30 min at room temperature with anti-mouse CD3-APC-Cy7, CD3-AF488, CD4-BV786, CD8-PE-Cy7(Tonbo), CD8-PE-Cy5, CD44 PerP-Cy5.5, CD44 PE-Cy7, CD62L-FITC, CXCR3-BV-650, CXCR3-PE-Dazzle; For monocytes, cells were stained with Cd11b APC/Cy7, Ly6C BV711, Ly6G FITC and CCR2 PE/Cy7. All the antibodies were purchased from Biolegend unless specified.

### Intracellular staining

For intracellular cytokine staining, one million of lung single cells were Fc blocked with anti CD16/32 and live/dead stained with ghost dye BV510 (Tonbo), and surface stained as described above. The cells were fixed, permeabilized with CytoFix/CytoPerm and stained with anti-mouse granzyme B-conjugated BV421, and APC conjugated Perforin. For interferon gamma detection (IFNγ), lung single cells prepared from mock and IAV infected mice were stimulated with 10uM IAV peptide NP_366-374_ for 5 hours with Brefeldin A, in RPMI medium containing 10% FBS and supplemented with antibiotics, prior to intracellular staining with anti-mouse IFNγ-PE-Dazzle 594 (*26*). A BD FACS Symphony or SONY MA900 flow cytometers were used to acquire 100,000 events and data were analyzed using FlowJo (Tree Star). The list of the antibodies used in this study including their clone and catalogue number is provided in supplemental data (table S1)

### Immunofluorescence staining

Formalin-fixed and paraffin-embedded lung sections from mock and IAV-infected mice were prepared as previously described (*47*) and probed for epithelial or endothelial cells detection using monoclonal mouse anti-αSMA (1:10,000; A2547; Sigma-Aldrich, Darmstadt, Germany), rabbit anti-mouse CD31 (1:50; ab28364, Abcam, Cambridge, UK), or rabbit anti-mouse EpCAM (1:50, ab71916, Abcam, Cambridge, UK) antibodies. Tissues were incubated with corresponding secondary antibodies goat anti-mouse IgG2a-AF488 (1:200; A-21131, ThermoFisher), anti-rabbit IgG-AF546 (1:200; Invitrogen, Carlsbad, CA), in 7% goat serum/PBS for 1h at RT, nuclei were counterstained with DAPI. All the images were acquired using a Leica DMi8 Thunder Imager fluorescent microscope and analyzed using Image J software.

### *In vivo* CXCR3 neutralization

IAV-Infected WT mice were intraperitoneally injected with 250ug (200uL) of anti-mouse CXCR3 monoclonal antibody (WT-PR8+αCXCR3) (BioXCell, clone CXCR3-173) (*45*) every alternate day starting day 4 post IAV infection. Mice were euthanized on days 7 and 14 post-infection for further investigations.

### IAV titers

The lung IAV load was determined *via* endpoint dilution assay and expressed as 50% tissue culture-infective dose (TCID_50_). The partial lobe of right lung from mock or IAV-infected mice were homogenized in PBS with volume normalized to the lung weight and stored at -80C until use. The 10-fold dilutions of lungs lysate supernatants were mixed with 3×10^3^ Madin-Darby canine kidney cells (MDCK), in 4 replicates, in DMEM containing 0.0002% L-1-(tosylamido-2-phenyl) ethyl chloromethyl ketone (TPCK)-treated trypsin (Worthington Biochemical) with antibiotics. After 6 days of incubation (370C with 5% CO_2)_, the cells were fixed with formalin and stained with 0.3% crystal violet solution. For each animal, viral titers were obtained using serial dilutions on MDCK monolayers and normalized to the total volume of lung homogenate supernatant (*48*).

### Single-cell RNA-seq library preparation and sequencing

From single-cell suspensions of mock and IAV-infected lungs at 7- and 14-days pi, CD8^+^ T cells were FACS sorted by positive staining with anti-mouse CD8-APC and CD3-FITC using BD Aria II flow cytometer sorter. 2×10^6^ cells were resuspended in media containing 10% DMSO and 20% FBS and allowed to slow freeze in Mr. Frosty. 3’ single-cell gene expression libraries (v3.0) were constructed using the 10x Genomics Chromium system. Single-cell library preparation was done by Singulomics Corporation (https://singulomics.com/). A pair-end 150 bp sequencing was performed to produce high-quantity data on an Illumina HiSeq 2000 platform (Illumina, San Diego, California, USA).

### Data alignment and sample aggregating

Sample demultiplexing, barcode processing, and unique molecular identifier (UMI) counting were performed by the Cell Ranger v5.0 (*49*). Briefly, fastq files were first extracted from the raw bcl files with the cellranger *mkfastq* demultiplexing pipeline. Then the fastq files were mapped to the mouse reference genome (mm10) to generate the count matrices by using the “cellranger count” pipeline with the default filtering parameters. Finally, the count matrices of all three time points were loaded into R 4.0.2 with the Seurat package (version 3.2.2) (*50*). To obtain high-quality data for the downstream analysis, we discard cells with less than 200 and more than 6,000 expressed genes, or the fraction of transcripts mapped to mitochondrial genes larger than 1%. The expression level of each gene was normalized by using *NormalizeData* function of Seurat. Finally, the datasets were integrated using the Seurat integration workflow.

### Dimensionality reduction and clustering

Principal component analysis (PCA) was performed to identify the cell clusters based on the highly variable genes. The principal component (PC) number where the percent change in variation between the consecutive PCs with less than 0.1% was selected as the optimal PC number. To create the Uniform Manifold Approximation and Projection (UMAP), the graph-based clustering was performed on the PCA-reduced data for clustering analysis (resolution=0.8) with *FindClusters* function.

### Cell type identification and differential expression analysis

To determine the cell type of each cluster, the gene markers for each cluster were identified by using the *FindAllMarkers* function in Seurat. Then, the cell type for each cluster was assigned by using the *CellMarker* and *PanglaoDB* database as the reference based on marker genes, with manual correction. Differentially expressed genes (DEGs; adjusted P-value < 0.05 and the average |log2FC| > 0.25) were identified by MAST (*51*) across experimental groups. Gene Set Enrichment Analysis (GSEA) of Kyoto Encyclopedia of Genes and Genomes (KEGG) were performed by using richR package (https://github.com/hurlab/richR) and significant pathways were identified with the P-value <0.05.

### Gene set variation analysis (GSVA)

Pathway activities in individual cells were assessed using the GSVA package (version 1.40.1) (*52*) with standard settings based on the KEGG dataset. To assess differential activities of pathways between different types of cells, we contrasted the activity scores for CD8^+^ T_eff_ against the CD8^+^ T_N_ and CD8^+^ T_CM_ cells by using the Wilcoxon test, and an adjusted P-value < 0.05 was used as the cutoff value for significant pathways identification.

### Cell-cell communication

To determine global communications among cells, CellChat (*53*) was used for the cell-cell communication analysis. The gene expression data of cells were used as the input and then the CellChat object was created with the metadata, including the cell type and group information. Then, the significant ligand-receptor pairs among cell groups were identified by performing a permutation test and categorized into signaling pathways. Pathways were filtered out if there are only 10 cells in certain cell groups. Next, the key incoming and outgoing signals were predicted for each cell group as well as global communication patterns by leveraging pattern recognition approaches. To cluster the signaling pathways, the similarity of pathways was measured and performing manifold learning from both functional and topological perspectives. The overall communication probability analysis was performed across all the datasets.

### Transcription factor (TF) regulons prediction

Gene regulatory network (GRN) was generated using the SCENIC package (*54*). Briefly, the raw expression matrix for the cells of all samples was filtered by keeping genes with default paramaters and then the co-expressed gene modules and the potential TF targets (regulons) for each module were identified with the GENIE3 based on the expression matrix. Cis-regulatory motif analysis was performed by scanning two TFs databases (https://resources.aertslab.org/cistarget/) (*mm10 refseqr80 10kb_up_and_down_tss.mc9nr.feather*, *mm10 refseqr80 500bp_up_and_100bp_down_tss.mc9nr.feather*) with the *RcisTarget* implemented in SCENIC. The modules with significant motif enrichment were kept and termed as regulons. To visualize regulon activity for each cell, the Area Under the Curve (AUC) scores (regulon activities) in each cell were computed and binarized the regulon network activity based on the AUCell algorithm, and we used binary regulon activity matrix to visualize regulon activity.

### Statistical analysis

Statistical analysis was performed using Prism 9 (GraphPad Software), and significance was determined by unpaired two-tail T-test for the comparison of means between two groups, one-way ANOVA with Tukey post hoc test was used for multiple comparisons between groups, two-way ANOVA test was used for flow cytometric kinetic data and body weight change data with only the significance of comparisons by timepoint are shown in the figures for clarity. P-values less than 0.05 were considered significant. All the experiments were repeated at least twice.

## Acknowledgements

We thank Jay Kolls (Center for Translational Research in Infection and Inflammation Tulane School of Medicine) for his valuable inputs in scRNAseq and overall interpretation of data. We thank Michaela Lano (UND school of Medicine and Health Sciences) and Shahram Solaymani-Mohammadi (UND school of Medicine and Health Sciences) for their critical reading of the manuscript.

## Author contributions

N.K. conceived the research and designed the experiments. D.Y., T.S. and J.T. performed and analyzed the experiments. K.G and N.K analyzed and interpreted the scRNAseq data. N.K., K.G., D.Y., M.O., J.X., J.H., J.S. and Z.W. wrote and revised the manuscript. All authors provided comments to the manuscript. All authors have seen and approved the manuscript, which has not been accepted or published elsewhere.

## Data and materials availability

All data associated with this study are present in the paper or the Supplementary Materials. The scRNAseq data reported in this study have been deposited in the NCBI Gene Expression Omnibus (GEO; www.ncbi.nlm.nih.gov/geo/) under the accession number GSE186839. The scRNA-seq datasets for whole lung were collected from NCBI’s BioProject database (PRJNA733762). The script for the preprocessing of the data is publicly available (free) at Github (https://github.com/guokai8/Cd8_T_cell) and deposited in the Synapse (ID: syn26410284).

---

**Fig. S1.**
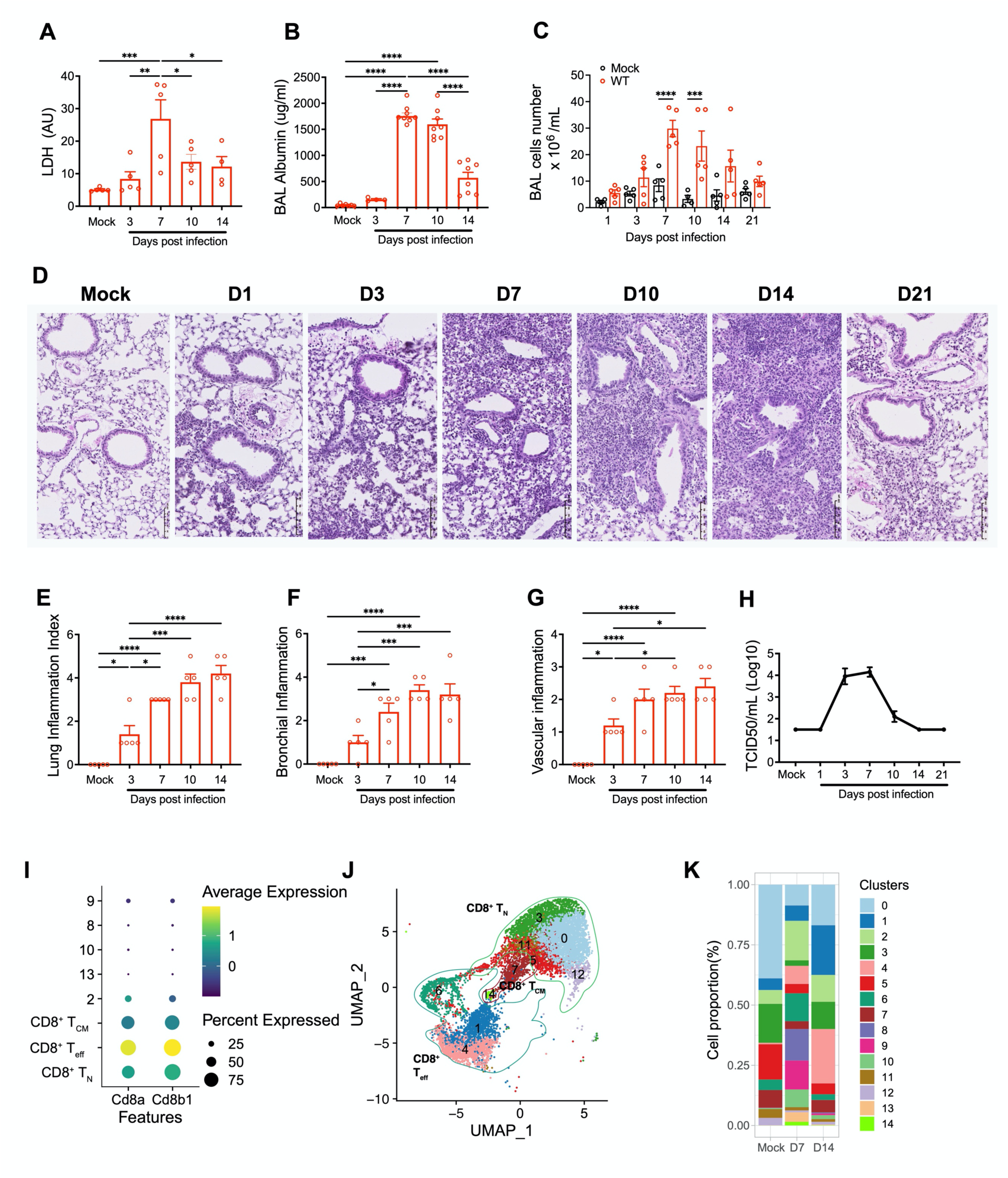
The kinetics of lung inflammation and pathology in influenza model. WT (C57BL/6) mice were mock-infected with PBS (mock) or infected with 1,000 PFUs of IAV (PR8) intranasally. At indicated time points, mice were euthanized, the BAL fluid and lungs were aseptically collected. The FACS sorted CD8^+^ T cells from mock and IAV-infected mice were subjected to scRNAseq analysis and lung single cells were used for flow cytometry characterization of CD8^+^ T cells. **(A-C).** The kinetics of of LDH **(A)**, albumin **(B)**, and total number of leukocytes **(C)** in the BALF from mock and PR8-infected mice. Data shown as means ± SEM with individual data shown as dot. Data is representatice of two independent experiments with n=5 per group and time point. one-way ANOVA with Tukey post hoc test was used in A-B and two-way ANOVA, with Dunn-Sidak multiple comparisons correction between mock and infected mice at each time point in **(C)**, and **(D)** H&E staining of paraffin-embedded lung sections from mock and PR8-infected mice (WT-PR8) at different time points. Data is representative of 3 different experiments with n=5 per group, scale bar=100μm, images were taken at 20X magnification. **(E-G)** The inflammation and pathology in H&E-stained lung sections. Lung inflammation score index **(E)**, bronchial **(F)** and vascular/endothelial **(G)** damage scores. Non-parametric Kruskal-Wallis test was used for multiple comparisons of median between mock and PR8-infecetd for statistical significance, representative of 2 different experiments, n=5. * p< 0.05, *** p< 0.005 and **** p<0.001. **(H)** TCID_50_ Kinetics of influenza load in lung homogenates of PR8-infected mice. Data shown as means ± SEM, representative of 2 different experiments with n=5/group and time point. **(I)** Dot plot showing the expression of CD8a and CD8b in CD8^+^ subsets and clusters of total CD8^+^ T cells. **(J)** UMAP of CD8^+^ T cells transcriptomes after removal of clusters with low expression of CD8 markers (clusters 2, 8, 9, 10, and 13). **(K)** Distribution of CD8^+^ T cells clusters assigned by unsupervised analysis. Color indicates the clusters.

**Fig. S2.**
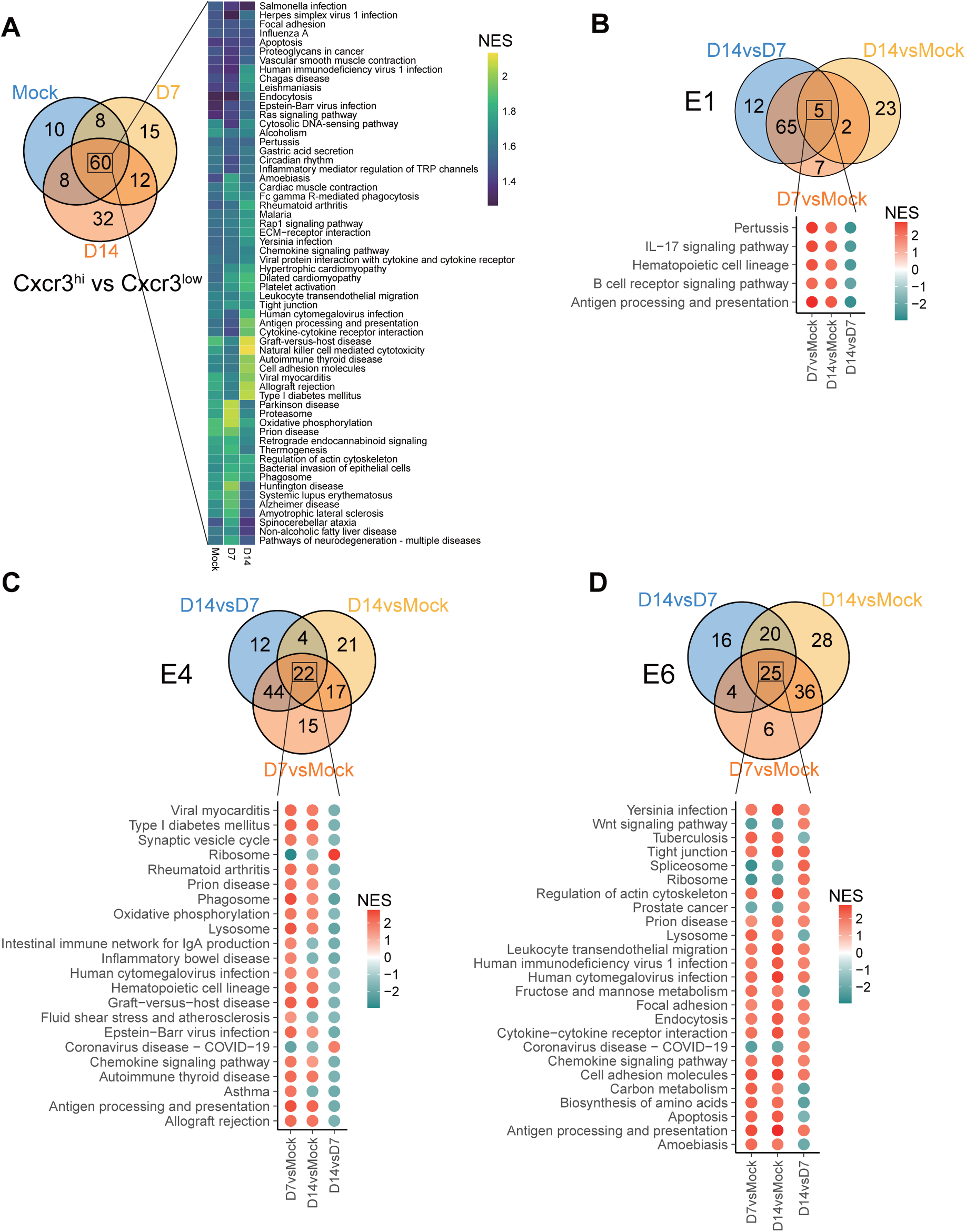
Landscape of significantly enriched pathways in of CD8^+^ T_eff_ cells. **(A)** Left: Venn Diagram showing the shared and unique significant pathways between Cxcr3^hi^ and Cxcr3^low^ CD8^+^ Teff clusters in mock and PR8 infecetd mice at D7 and D14 from Gene Set Enrichment Analysis (GSEA); Right: Heatmap shows the shared significant pathways between Cxcr3^hi^ and Cxcr3^low^ CD8^+^ Teff clusters in mock and PR8 infecetd mice at D7 and D14.The color stands for the normalized enrichment score (NES) value. **(B)** Top: Venn Diagram showing the shared and unique significant KEGG pathways between Cxcr3^hi^ and Cxcr3^low^ CD8^+^ Teff clusters in mock and PR8 infecetd mice at D7 versus Mock, D14 versus Mock, and D14 versus D7 of E1 from GSEA; Bottom: Dot plot shows the shared significant pathways among all three comparisons. The color stands for (up-regulated (red) or down-regulated (blue) in the corresponding groups. **(C)** Top: Venn Diagram showing the shared and unique significant KEGG pathways between Cxcr3^hi^ and Cxcr3^low^ CD8^+^ Teff clusters in mock and PR8 infecetd mice at D7 versus Mock, D14 versus Mock, and D14 versus D7 of E4 from GSEA; Bottom: Dot plot shows the shared significant pathways among all three comparisons. The color stands for (up-regulated (red) or down-regulated (blue) in the corresponding groups). **(D)** Top: Venn Diagram shows the shared and unique significant KEGG pathways between Cxcr3^hi^ and Cxcr3^low^ CD8^+^ Teff clusters in mock and PR8 infecetd mice at D7 versus Mock, D14 versus Mock, and D14 versus D7 ofof E6 from GSEA; Bottom: Dot plot shows the shared significant pathways among all three comparisons. The color stands for the normalized enrichment score (NES) value (up-regulated (red) or down-regulated (blue) in the corresponding groups).

**Fig. S3.**
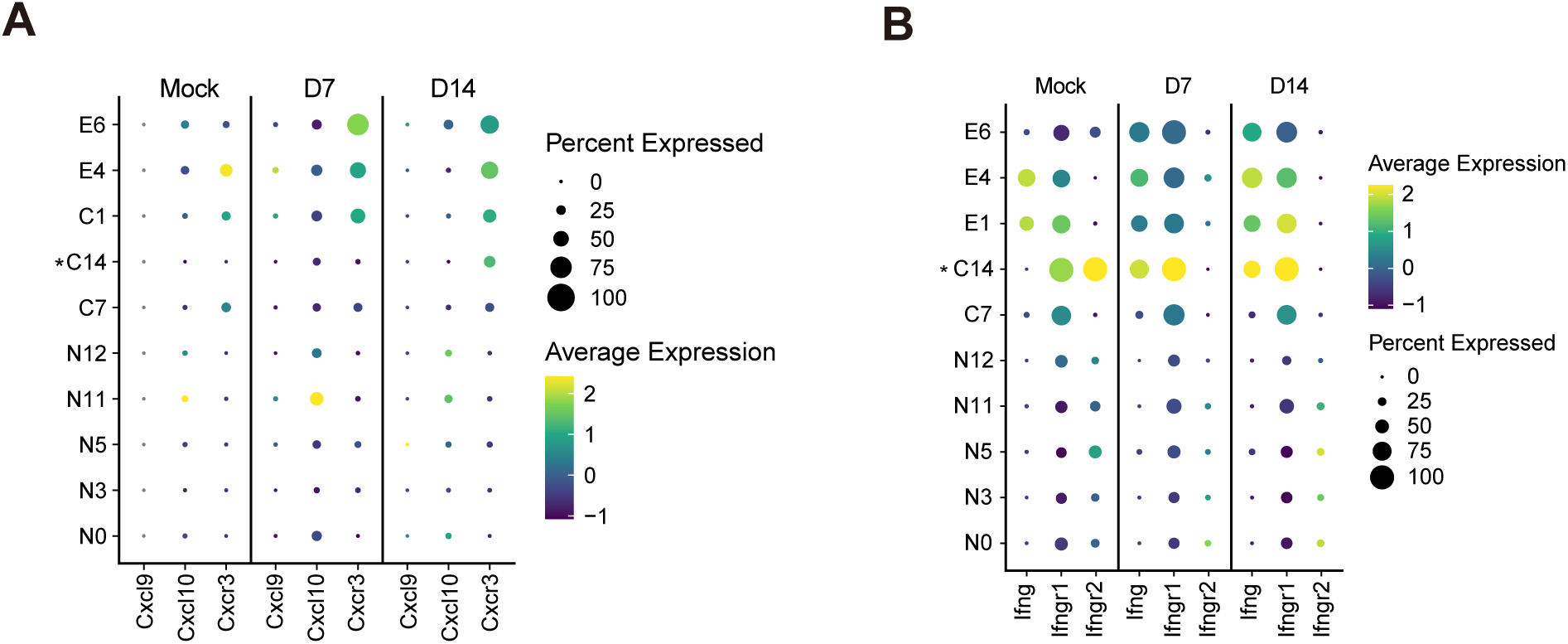
Landscape of cell-cell communications between CXCR3^+^ CD8^+^ T cells in mock and PR8-infected mice. **(A)** Dot plot representing ligand and receptors expression levels of CXCL signaling pathway for 10 clusters generated. * Indicate that C14 had less than 10 cells at one of the time points. **(B)** Dot plot representing ligand and receptors expression levels of IFN-II signaling pathway for 10 clusters generated. * Indicate that C14 had less than 10 cells at one of the time points.

**Fig. S4.**
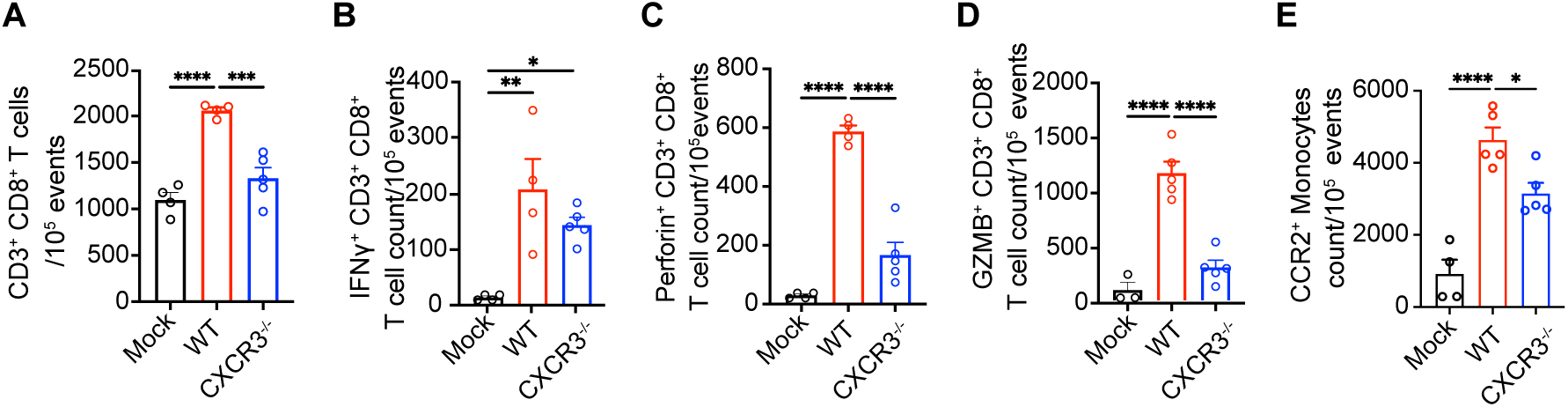
A comparison of Lung inflammation and pathology in influenza-infected WT and CXCR3^-/-^ mice. WT and CXCR3^-/-^ mice were mock-infected with PBS (mock) or infected with 1,000 PFUs of IAV (PR8) intranasally. At 7 dpi mice were euthanized, and lungs aseptically collected for further analysis. The flow cytometry immunohneotyping was performed in single-cell digests of lungs from WT and CXCR3^-/-^ mice. **(A-D)** The frequencies of total **(A)**, IFN-γ^+^ **(B)**, perforin^+^ **(C)**, GZMB^+^ **(D)** CD8^+^ T cells between PR8 infected WT and CXCR3^-/-^ mice. **(E)** The frequency of CCR2^+^ monocytes between PR8 infected WT and CXCR3^-/-^ mice. One-way ANOVA with Turkey post hoc test was used for multiple comparisons of the means between groups. The data is representative of one experiment, n=5/group. * p< 0.05, ** p< 0.01, *** p< 0.005 and **** p< 0.001.

**Fig. S5.**
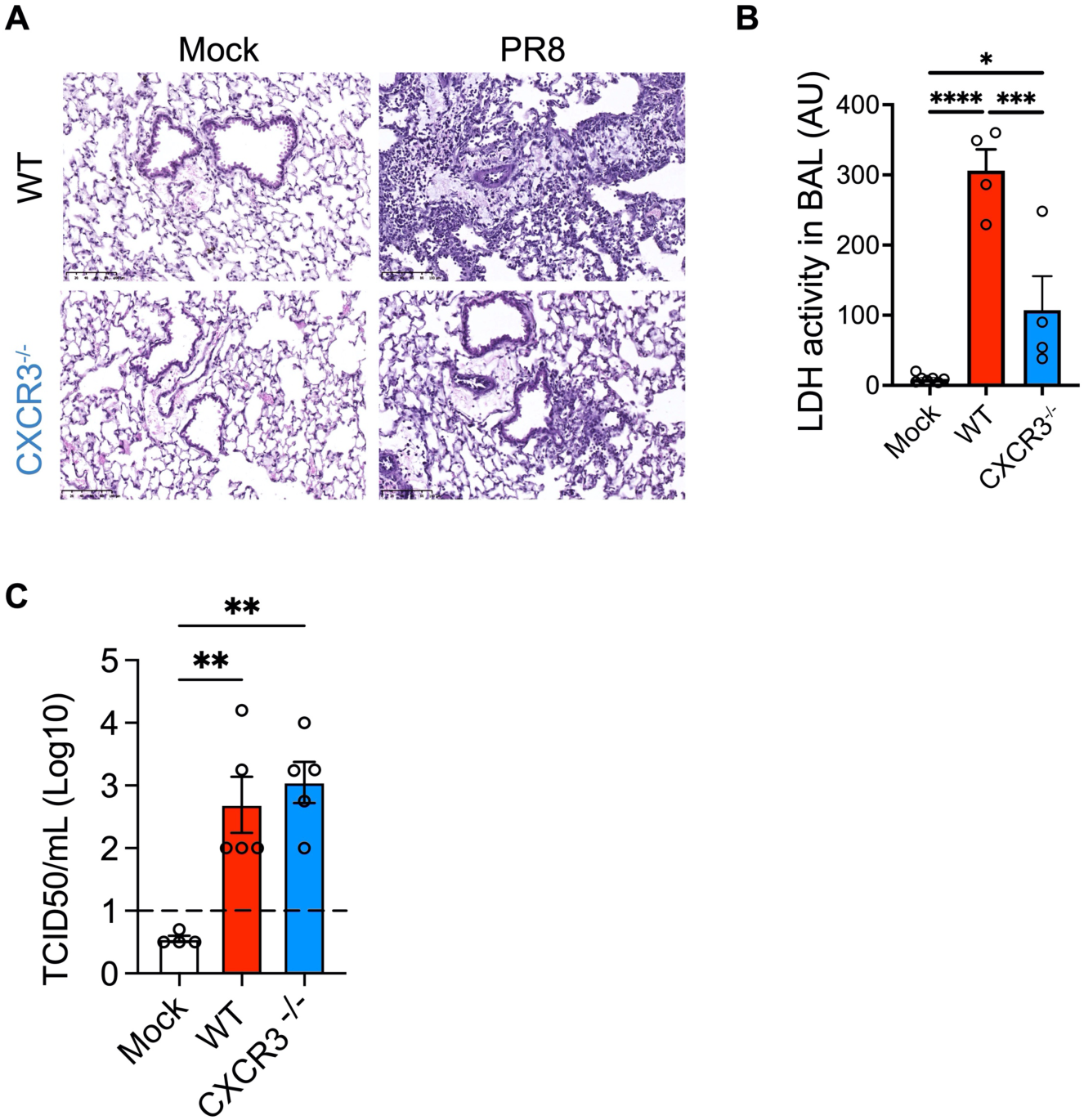
A comparison of Lung inflammation and pathology in influenza-infected WT and CXCR3^-/-^ mice. WT and CXCR3^-/-^ mice were mock-infected with PBS (mock) or infected with 1,000 PFUs of IAV (PR8) intranasally. At 7 dpi mice were euthanized, and lungs and BALF were aseptically collected for further analysis. **(A)** H&E staining of paraffin-embedded lungs sections from WT (top) mice and CXCR3^-/-^ mice (bottom) at 7 dpi. Representative images of mock (left) and PR8-infected (right) mice with n=5 per group, images taken at X20 magnification with scale bar 100um. **(B)** Level of LDH in BAL of PR8 infceted WT and CXCR3^-/-^ mice. Data shown as means ± SEM. One-was ANOVA with Tukey post hoc test was used for multiple comparisons of means between groups. * p< 0.05, *** p< 0.005 and **** p<0 .001 **(C)** TCID_50_ influenza viral load in lung homogenates of PR8-infected WT and CXCR3^-/-^ mice. Data shown as means ± SEM. One-way ANOVA test for multiple comparisons of means between groups was used. * p< 0.05, *** p< 0.005 and **** p<0 .001

**Table S1.** Antibodies are used for Flow cytometry.

| **Antibody** | **Dilution** | **Clone** | **Catalog#** | **Company** |
| --- | --- | --- | --- | --- |
| CD3<br>APC/Cy7 | 1:200 | 145-2C11 | 100330 | Biolegend |
| CD4 BV711 | 1:200 | RM4-5 | 100550 | Biolegend |
| CD8 PE/Cy7 | 1:200 | 2.43 | 60-1886-U100 | TONBO biosciences |
| CD62L<br>FITC | 1:200 | MEL-14 | 104406 | Biolegend |
| CD44 Percp<br>Cy5 | 1:200 | IM7 | 103032 | Biolegend |
| CD44<br>PECy5 | 1:200 | IM7 | 103009 | Biolegend |
| CXCR3<br>BV650 | 1:100 | CXCR3-173 | 126531 | Biolegend |
| CXCR3 PE-Dazzle | 1:100 | CXCR3-173 | 126534 | Biolegend |
| IFNγ<br>PE/dazzle | 1:50 | XMG1.2 | 505846 | Biolegend |
| Perforin<br>APC | 1:75 | S16009A | 154304 | Biolegend |
| Granzyme<br>B. BV421 | 1:75 | QA18A28 | 396414 | Biolegend |
| Ghost Dye-BV 510 | 1:500 |  | 13-0870 | TONBO biosciences |
| anti-mouse<br>CD16/CD32 | 1:200 | 2.4G2 | 553142 | BD Biosciences |
| CD11b<br>APC/Cy7 | 1:200 | M1/70 | 101226 | Biolegend |
| Ly6C<br>BV711 | 1:400 | HK1.4 | 128037 | Biolegend |
| Ly6G FITC | 1:200 | 1A8 | 127605 | Biolegend |
| CCR2<br>PE/Cy7 | 1:100 | SA203G11 | 150612 | Biolegend |

**Data S1. Wilcoxon test of pathways activities for CD8+ Teff against CD8+ TN and CD8+ TCM**

**Data S2. Shared or unique DEG subsets between Cxcr3hi and Cxcr3low at mock, 7 and 14 dpi.**

Shared: DEGs are shared among mock, 7 and 14 dpi; D7_Mock: DEGs are shared at mock and 7 dpi; D14_Mock: DEGs are shared at mock and 14 dpi; D14_D7: DEGs are shared at 7 and 14 dpi.

**Data S3. Shared or unique significant pathway subsets between Cxcr3hi and Cxcr3low at mock, 7 and 14 dpi.**

Shared: significant pathways are shared among mock, 7 and 14 dpi; D7_Mock: significant pathways are shared at mock and 7 dpi; D14_Mock: significant pathways are shared at mock and 14 dpi; D14_D7: significant pathways are shared at 7 and 14 dpi.

**Data S4. Shared or unique DEG subsets among D7 vs Mock, D14 vs Mock and D14 vs D7 in cluster E1.**

Shared: DEGs are shared among all three comparisons; D14vsMock_D14vsD7: DEGs are shared at mock and 7 dpi; D14_Mock: DEGs are shared at mock and 14 dpi; D14_D7: DEGs are shared at 7 and 14 dpi.

**Data S5. Shared or unique significant pathway subsets among D7 vs Mock, D14 vs Mock and D14 vs D7 in cluster E1.**

Shared: significant pathways are shared among all three comparisons; D14vsMock_D14vsD7: significant pathways are shared at mock and 7 dpi; D14_Mock: significant pathways are shared at mock and 14 dpi; D14_D7: significant pathways are shared at 7 and 14 dpi.

**Data S6. Shared or unique DEG subsets among D7 vs Mock, D14 vs Mock and D14 vs D7 in cluster E4.**

Shared: DEGs are shared among all three comparisons; D14vsMock_D14vsD7: DEGs are shared at mock and 7 dpi; D14_Mock: DEGs are shared at mock and 14 dpi; D14_D7: DEGs are shared at 7 and 14 dpi.

**Data S7. Shared or unique significant pathway subsets among D7 vs Mock, D14 vs Mock and D14 vs D7 in cluster E4.**

Shared: significant pathways are shared among all three comparisons; D14vsMock_D14vsD7: significant pathways are shared at mock and 7 dpi; D14_Mock: significant pathways are shared at mock and 14 dpi; D14_D7: significant pathways are shared at 7 and 14 dpi.

**Data S8. Shared or unique DEG subsets among D7 vs Mock, D14 vs Mock and D14 vs D7 in cluster E6.**

Shared: DEGs are shared among all three comparisons; D14vsMock_D14vsD7: DEGs are shared at mock and 7 dpi; D14_Mock: DEGs are shared at mock and 14 dpi; D14_D7: DEGs are shared at 7 and 14 dpi.

**Data S9. Shared or unique significant pathway subsets among D7 vs Mock, D14 vs Mock and D14 vs D7 in cluster E6.**

Shared: significant pathways are shared among all three comparisons; D14vsMock_D14vsD7: significant pathways are shared at mock and 7 dpi; D14_Mock: significant pathways are shared at mock and 14 dpi; D14_D7: significant pathways are shared at 7 and 14 dpi.

